# Evolution of binding preferences among whole-genome duplicated transcription factors

**DOI:** 10.1101/2021.07.27.453962

**Authors:** Tamar Gera, Felix Jonas, Roye More, Naama Barkai

**Affiliations:** Department of Molecular Genetics, Weizmann Institute of Science; Rehovot 76100, Israel

## Abstract

Throughout evolution, new transcription factors (TFs) emerge by gene duplication, promoting growth and rewiring of transcriptional networks. How TF duplicates diverge is known for only a few studied cases. To provide a genome-scale view, we considered the 35% of budding yeast TFs, classified as whole-genome duplication (WGD)-retained paralogs. Using high-resolution profiling, we find that ~60% of paralogs evolved differential binding preferences. We show that this divergence results primarily from variations outside the DNA binding domains (DBDs), while DBD preferences remain largely conserved. Analysis of non-WGD orthologs revealed that ancestral preferences are unevenly split between duplicates, while new targets are acquired preferentially by the least conserved paralog (biased sub/neo-functionalization). Dimer-forming paralogs evolved mostly one-sided dependency, while other paralogs interacted through low-magnitude DNA-binding competition that minimized paralog interference. We discuss the implications of our findings for the evolutionary design of transcriptional networks.

## Main

Transcription factors (TFs) bind at regulatory regions to activate or repress transcription. Cells express hundreds of TFs that together transit cells between different expression programs. Despite rapid advances, our understanding of transcriptional networks is still fragmented (*1*). For example, different TFs that bind to similar DNA sequences *in-vitro* (*2–10*) localize to different genomic sites *in-vivo* through mechanisms still poorly understood. Further, with increasing organism complexity, new TFs emerge, yet we know little about the means through which these emerging TFs adapt new targets and integrate into the existing transcriptional network.

Gene duplication is the major source of new TFs (*11–15*), with whole-genome duplications (WGDs) playing a particularly important role (*16–26*). In budding yeast, ~35% of all TFs are associated with a single WGD event dating ~100 million years ago (Fig. 1A) (*19, 20*). We reasoned that this set of TF duplicates, all generated at the same time and subjected to the same evolutionary history, provides a convenient platform for studying the fate of retained duplicated TF genes.

**Fig. 1.**
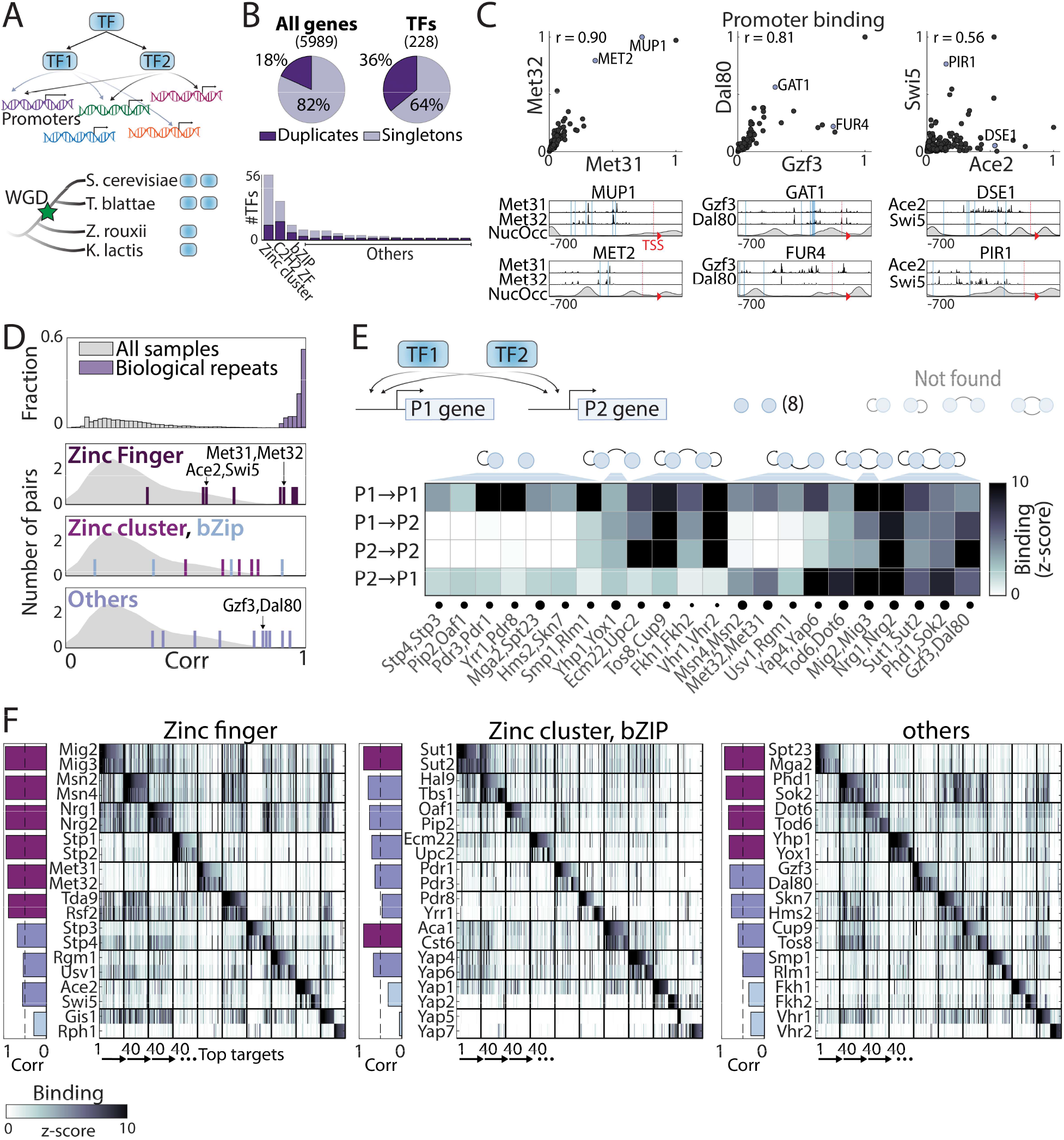
Promoter binding preferences of WGD TF-duplicates. **(A-B)** Whole-genome duplication (WGD) shaped the budding yeast transcription network: (A) TF duplicates (paralogs) can diverge to bind different targets. (B) In *S. cerevisiae*, ~35% (*82*) of all present day TFs are retained WGD paralogs (see fig. S1). **(C-D)** Binding profiles of TF paralogs: Shown in (C) are promoter binding preferences of the indicated TFs (top, each dot is a promoter, r: Pearson’s correlation). Bottom plots show binding signal over individual promoters, with lines indicating transcription start site (TSS, red dashed) and locations of *in-vitro* motifs (blue). Nucleosome positioning shown in grey. (D) Distributions of correlations between promoter binding preferences of different TFs (grey), between biological repeats (purple, top) and between paralogs (colored lines, bottom). **(E)** Auto- and cross-promoter binding by TF paralogs: Promoter binding signal is shown as z-score, with potentially formed circuits shown on top. Note that 22/30 pairs form six of nine possible circuits. **(F)** Binding to top-scoring promoters: The 40 top-bound promoters by each TF pair were selected (see table S2), ordered along the x-axis, and color-coded according to binding strength by each TF (y-axis). Bars on the left show Pearson’s correlations of binding preferences between the indicated pairs.

TF duplicates (paralogs) can diverge through changes in expression, regulation or function (*27–44*). Of particular interest is the selection of *in-vivo* binding sites, as these define potential regulatory targets. Mechanisms driving such divergence include changes in co-factor binding (*45*) or in DNA motif preferences (*46–51*). The prevalence of these different scenarios is still unclear, since studied cases considered only a few paralogs, of different ages and origins, and divergence was studied at individual targets. We therefore aimed to provide a genome-scale view, by comparing genome-wide binding preferences among the full set of WGD TFs in budding yeast.

## Results

### Divergence of binding preferences among WGD TFs

Within the *Saccharomyces cerevisiae* genome database (SGD) (*52*), 82 DNA binding domain (DBD)-containing proteins are classified as WGD-retained paralogs (Fig. 1A,B). We refined this list to include only pairs where both paralogs act as specific TFs (35 pairs, table S1), and defined the binding locations of these TFs across the genome using ChEC-seq (*53*). A total of 30 pairs (60 TFs) were successfully profiled, as verified by data reproducibility (Pearson’s r>0.95 in promoter binding preferences, Fig. 1C,D) and manual literature survey (fig. S1). A large fraction of TFs were bound at their own or their paralog’s promoters, potentially forming regulatory circuits (*15*) (Fig. 1E).

Perhaps unexpectedly, binding preferences were conserved (Pearson’s r>0.8) among ~40% of paralogs, most notably these of the C2H2 zinc-finger family (6/10 pairs, e.g. Met31/Met32, Fig. 1C,D). Furthermore, most diverging paralogs still shared promoter targets (Fig. 1F). In some cases, the two duplicates localized to the same promoters but with different relative strengths (e.g. Gzf3/Dal80, Fig. 1C). In other cases, some promoters were bound by both paralogs, while others by just one (e.g. Ace2/Swi5, Fig. 1C). Therefore, binding preferences diverge at a rate that differs between pairs, and, within each pair, differs between individual promoters.

### Paralog diverge through variations outside their DBDs

TFs localize to genomic sites containing short motif sequences bound by their DBDs. The *in-vivo* binding could therefore diverge through DBD variations that modify motif preferences. To compare DBD sequences among paralogs, we aligned each pair and classified residues into these defining the DBD family (e.g. C/H residues in the C2H2 zinc-finger domains), those that contribute to DNA motif preference (*54*), and the remaining ones (Fig. 2A,B and fig. S2).

**Fig. 2.**
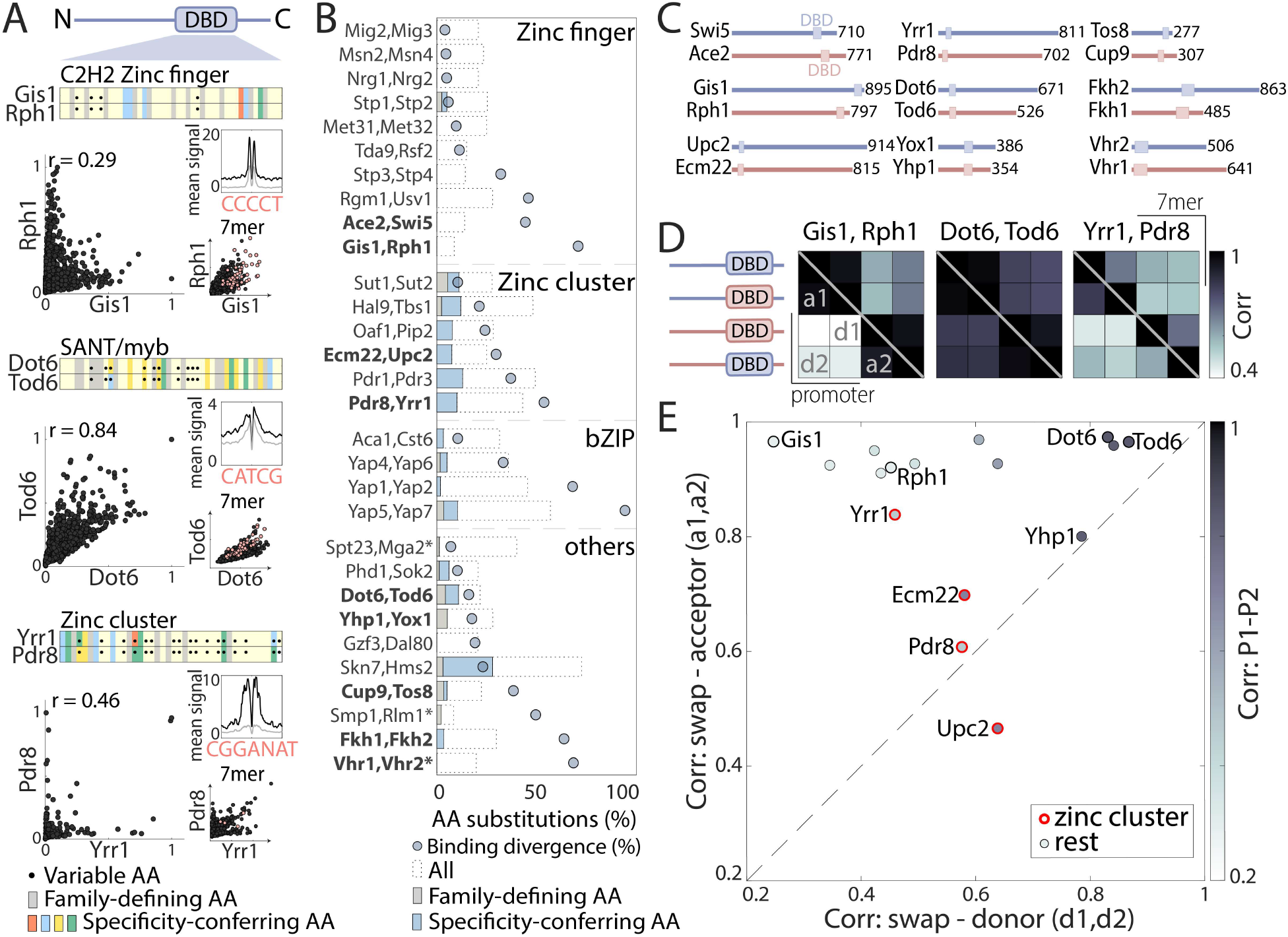
The contribution of DBDs to the divergence of TF paralogs. **(A-B)** Sequence variations between paralogs’ DBDs: For all pairs, Pfam-defined DBDs were aligned and residues classified into those conserved among all family members (grey) and specificity-conferring ones (colored; blue, red, yellow and green denoting basic, acidic hydrophobic and hydrophilic residues respectively, (*54*)), as shown in (A) for the three examples (black dots indicate positions varying between paralogs). The fraction and class of varying residues is summarized in (B) (*:specificity-conferring residues not defined). Note the little correspondence between DBD sequence variations and binding profile divergence (blue dots). Paralogs chosen for further analysis are highlighted in bold. Also shown in (A) are the average signal around *in-vitro*-motif occurrences (right, top), the respective binding signals across promoters (left) and at all 7-mer sequences (right bottom, red dots indicate 7-mers containing the *in-vitro* motif). **(C-E)** DBD swapping has a minor effect on binding preferences: DBDs of the indicated paralog pairs (C, protein length indicated) were swapped, and their binding profiles mapped. Shown in (D) are the correlations between binding preferences of the indicated TFs and their swapped variants (bottom triangle promoters, top triangles 7-mers; see also fig. S2). These correlations are summarized in (E) for all tested pairs (red indicates zinc-cluster TFs; shading depicts correlation between wild type paralogs). Note that outside the zinc-cluster family, DBD swapping is of little consequence for promoter binding preferences, even among highly divergent paralogs. Within the zinc-cluster family, DBD swapping affected binding profiles, but did not recover donor profiles.

Sequence conservation varied between DBD families (Fig. 2B). In particular, specificity-conferring residues often varied between paralogs of the fungal-specific zinc-cluster family (Fig. 2B), but remained invariant between paralogs of the C2H2 zinc-finger family and, to a lesser extent, in other families (e.g. Rph1/Gis1, Fig. 2A,B). Examining motif preferences derived from *in-vitro* data, we noted that reported preferences (*2, 54*) are often (although not always) similar amongst paralogs (Fig. 2A and fig. S2).

DBD variations may therefore contribute to the divergence of zinc-cluster paralogs, but appear to play a lesser role in paralogs of the C2H2 zinc finger and perhaps other families. To test this, we swapped DBDs between paralogs (Fig. 2C), reasoning that if DBD variations drive divergence, swapping DBDs would swap promoter preferences. Conversely, if critical variations are located outside the DBD, swapping will be of little effect. Consistent with our sequence analysis, DBD swapping perturbed binding for three of the four zinc-cluster TFs tested, although in none of these cases was DBD swapping sufficient for swapping promoter preferences (Fig. 2D,E). The fourth TF (Yrr1) remained largely invariant (Pearson’s r>0.8) to DBD swapping, as did the eleven additionally tested TFs, taken from six different families. Of note, this invariance to DBD swapping characterizing most TFs was observed not only when comparing promoter preferences, but also when comparing *in-vivo* preferences to DNA *7*-mers (Fig. 2D and fig. S2). We conclude that, for most duplicate pairs, the variations driving divergence in promoter binding preferences are located outside the DBDs.

### Dependencies and competitions between TF paralogs

Evolved interactions between paralogs could affect binding preferences. TFs that bind DNA as dimers, for example, may bind as heterodimers to a subset of sites. Paralogs may also compete for binding, either directly or by interacting with a shared co-factor. In the broader context, cooperative interactions, where a TF depends on its paralog, may increase mutation fragility whereas binding competition, which allows a TF to access its perturbed paralog’s sites, may increase mutation robustness (Fig. 3A). Both effects were observed in the context of proteinprotein interactions (*44*).

**Fig. 3.**
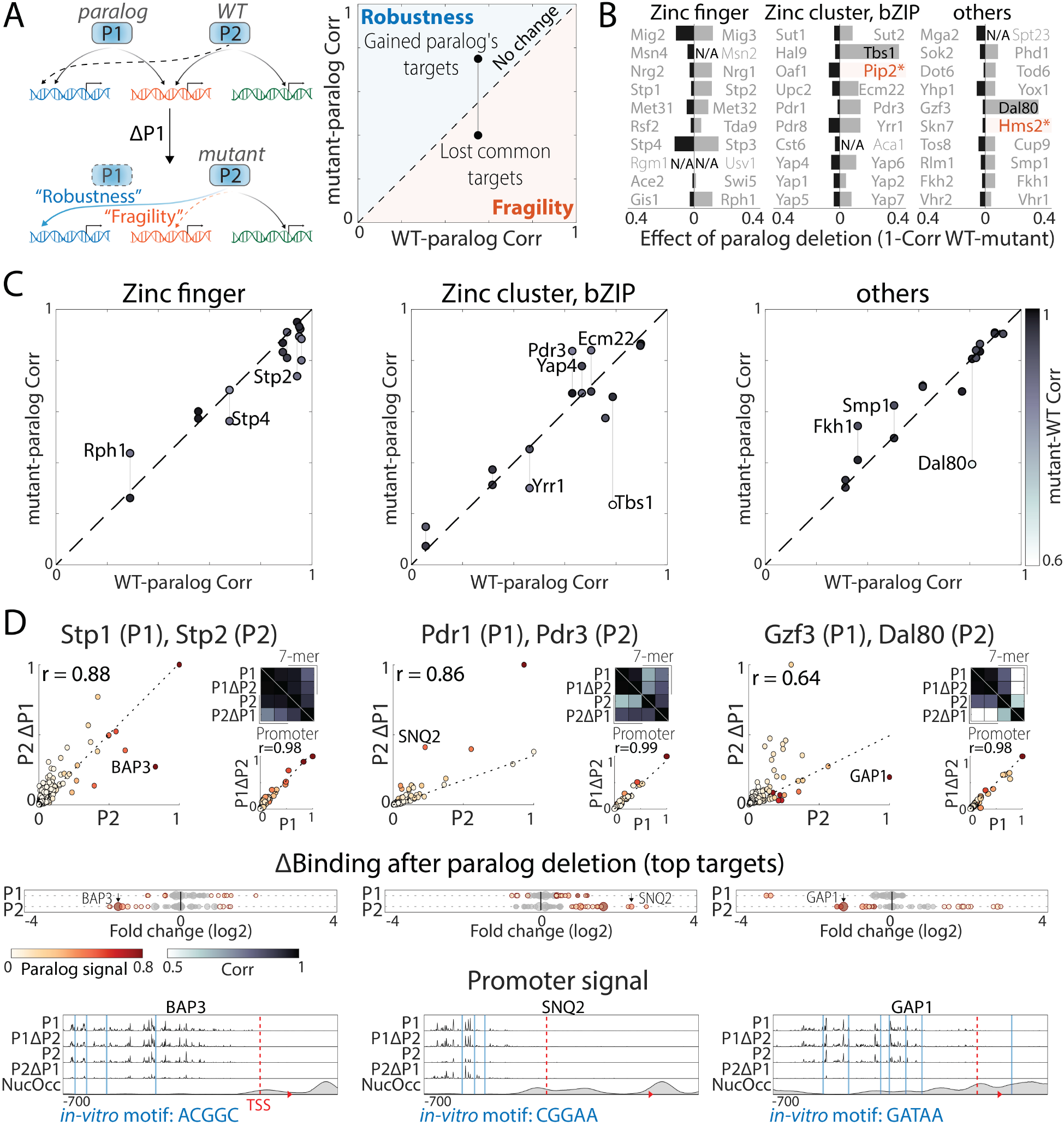
Interactions between TF paralogs may increase network fragility. **(A)** Paralogs’ contribution to mutation robustness or fragility: Following paralog deletion, a TF may gain access to its paralog’s unique sites, potentially compensating for the loss (“robustness”, blue line). Alternatively, paralogs may become interdependent (“Fragility”, orange line). These interactions can be visualized by comparing the TF’s binding preferences in wild type (x-axis) or paralog-deleted (y-axis) backgrounds to those of the paralog. **(B)** Paralog interactions are rare: the effect of paralog deletion on promoter binding preferences was measured for 55 TFs of 60 paralogs in our dataset. Shown is the effect of paralog deletion on binding preference for each TF. Note that most deletions were of little effect, or asymmetric in cases of large effects. Also indicated are substantial effects (TFs highlighted in black), and TFs that completely lost binding specificity (colored in orange, N/A: not profiled, see materials and methods). **(C-D)** Tendency for paralog interactions varies between TF families: (C) robustness/fragility analysis, as in (A) for all tested paralog pairs, divided into families. (D) For individual examples, binding is shown over all (top) or individual promoters (bottom), and as relative change following paralog deletion (middle; dot color and size indicate a TF’s and its paralog’s binding signal, respectively). Note that Stp2 and Dal80 lose binding to some of the targets shared with their respective paralog following paralog deletion (“fragility”), whereas Pdr3 gains binding to Pdr1 targets (SNQ2) upon the latter’s deletion (“robustness”).

To test the prevalence of cooperative or competitive interactions, we measured TF binding upon paralog deletion, testing 55/60 TFs in our dataset (Fig. 3B and fig. S3). Two TFs completely lost binding signals (Pip2, Hms2), and additional two lost binding to their respective paralogs’ targets (Dal80, Tbs1). These large-scale effects, however, were infrequent (Fig. 3B,C). Cooperative interactions were generally minor (e.g. Stp2), as were compensatory interactions (e.g. Pdr3 or Ecm22; Fig. 3C,D). Therefore, strong interactions between TFs paralogs are rare and existing ones tend to increase mutation fragility.

### Classifying paralogs’ evolutionary paths by analyzing non-WGD orthologs

Two prevailing models explain paralog divergence (*55–63*): In the first, ancestral functions split between duplicates (sub-functionalization) while, in the second, one duplicate retains ancestral functions, while the second adapts a new role (neo-functionalization, Fig. 4A). As a first approach to distinguish these two scenarios, we used phylogenetic sequence analysis to compare the evolutionary rates of the regions outside the conserved DBD (Fig. 4B). This analysis is informative, since a neo-functionalizing paralog undergoes a period of relaxed selection, followed by rapid evolution, and therefore evolves at an accelerated rate (*64, 65*). Diverging paralogs of the C2H2 zinc-finger family evolved symmetrically, at rates that did not distinguish between paralogs, consistent with sub-functionalization. By contrast, diverging paralogs of other families evolved asymmetrically, suggesting dominant neo-functionalization (Fig. 4B).

**Fig. 4.**
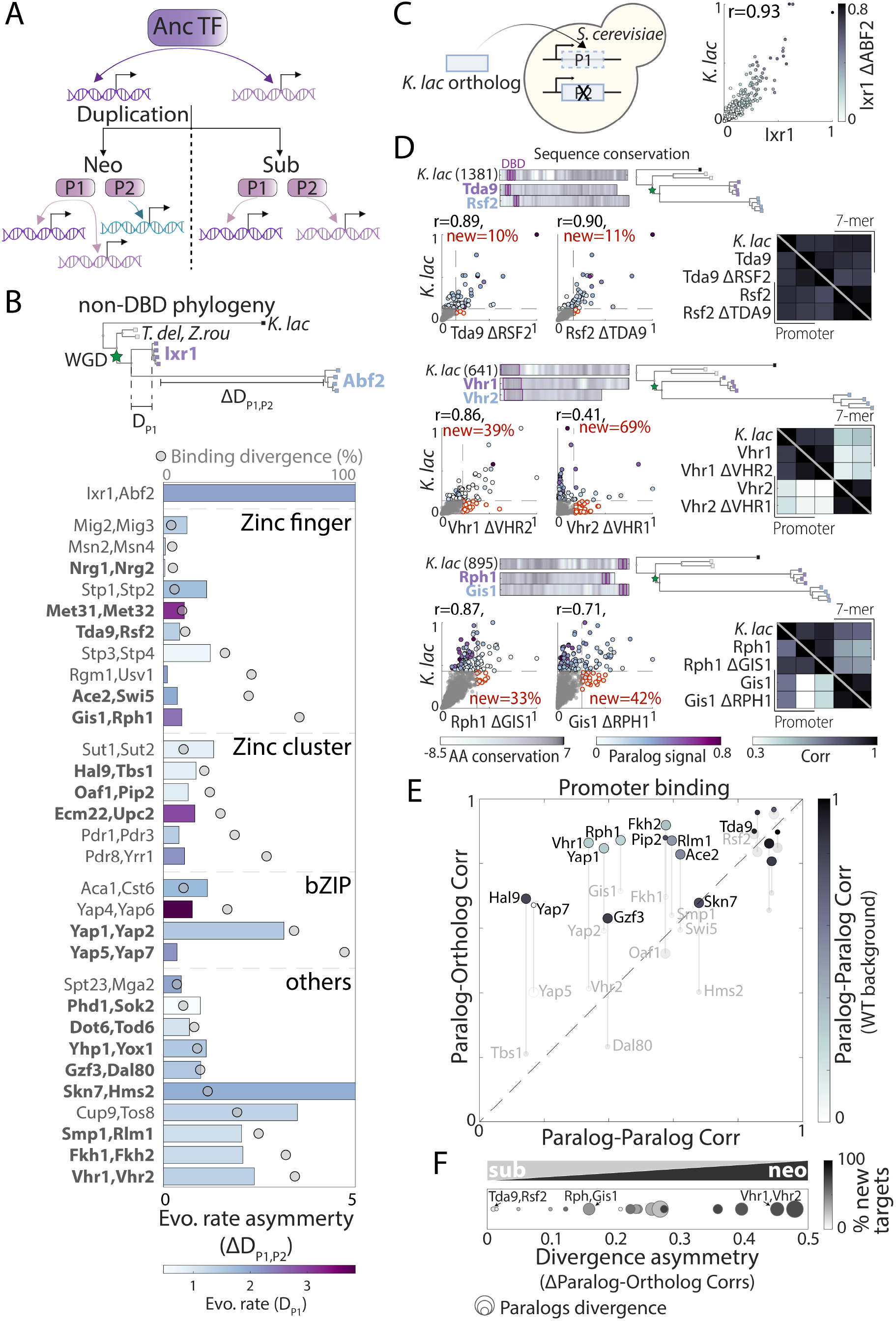
Classifying paralog divergence through comparison to non-WGD orthologs. **(A)** Models of evolutionary paths: Paralogs could diverge by acquiring new preferences (neo-functionalization) or by splitting ancestral preferences (sub-functionalization). **(B)** Sequence evolution: Top: Phylogenetic comparison of non-DBD sequences of the Ixr1/Abf2 paralogs, indicating distances from last common ancestor (LCA). Bottom: differences in sequence evolution rate between paralogs (evolutionary rate asymmetry, ΔD_P1,P2_, see methods for details). Color indicates rate of the conserved paralog (D_P1_), and grey dots indicate divergence of promoter binding preferences. Paralogs chosen for further analysis are highlighted in bold. **(C-F)** Binding preferences of non-WGD orthologs: Binding profiles of *K. lactis* orthologs were mapped within *S. cerevisiae* and compared to both paralogs (left). Ixr1 binding is tightly conserved between *S. cerevisiae* and *K. lactis* (right). Shown are the correlations of binding preference between the *K. lactis* and *S. cerevisiae* orthologs for specific examples (D) and for all tested paralogs (E; bigger spot size corresponds to the paralog with slower sequence evolution; see also fig. S4). Note the high similarity of binding preferences between each *K. lactis* TF with at least one of its *S. cerevisiae* paralogs, commonly the one experiencing slower sequence divergence. Divergence asymmetry is quantified in (F), with colors indicating the fraction of ‘new’ targets acquired by the less conserved paralog.

To test experimentally for sub- and neo-functionalization, we compared binding preferences of paralogs in our set to that of a corresponding non-WGD ortholog (*Kluyveromyces lactis* TF, expressed within *S. cerevisiae;* Fig. 4C). We reasoned that, in terms of binding preferences, this non-WGD ortholog might serve as a good proxy for the ancestor (*15, 66, 67*). This was the case in pairs with clear expectations: binding of the *K. lactis* ortholog of Ixr1/Abf2 was indistinguishable from that of Ixr1 (Fig. 4C), consistent with Abf2 adopting a mitochondria role (*68*) (note Abf2’s accelerated sequence evolution, Fig. 4B). Similarly, *K. lactis* orthologs of non-diverging duplicates retained highly similar binding preferences (Rsf2/Tda9 Fig. 4D and fig. S4 for additional pairs). Together, these results support the use of non-WGD orthologs to approximate ancestral preferences.

We next extended the analysis to divergent duplicates. We observed cases of clear sub- and neo-functionalization (Rph1/Gis1 and Vhr1/Vhr2, respectively Fig. 4D,E), but most pairs showed a combination of the two scenarios. Of note, in all 11 diverging cases, at least one paralog was more similar to the ortholog than to the other *S. cerevisiae* paralog and in 9 of them, the conserved paralog was also slower to evolve (Fig. 4E). Together, these results suggest that the prevalent route for diversification is through biased neo/sub-functionalization: ancestral preferences split unevenly between the duplicates, coupled to biased acquisition of novel targets (Fig. 4F and fig. S4).

### Divergence of zinc-cluster paralogs: motif preferences and paralog interactions

To gain molecular insights into the divergence of specific duplicates, we examined individual pairs, focusing first on the fungal-specific zinc-cluster family. TFs of this family bind DNA as dimers, recognizing a composite DNA motif that includes two spaced CGG sites (*69*). Binding specificity depends on the orientation of the DBD-bound CGG triplets, and the spacer length, which may depend on an unstructured linker flanking the DBD (Fig. 5A). As described above, when compared to other families, zinc-cluster paralogs showed a greater dependence on DBD variations (cf. Fig. 2) and were more likely to interact (cf. Fig. 3).

**Fig. 5.**
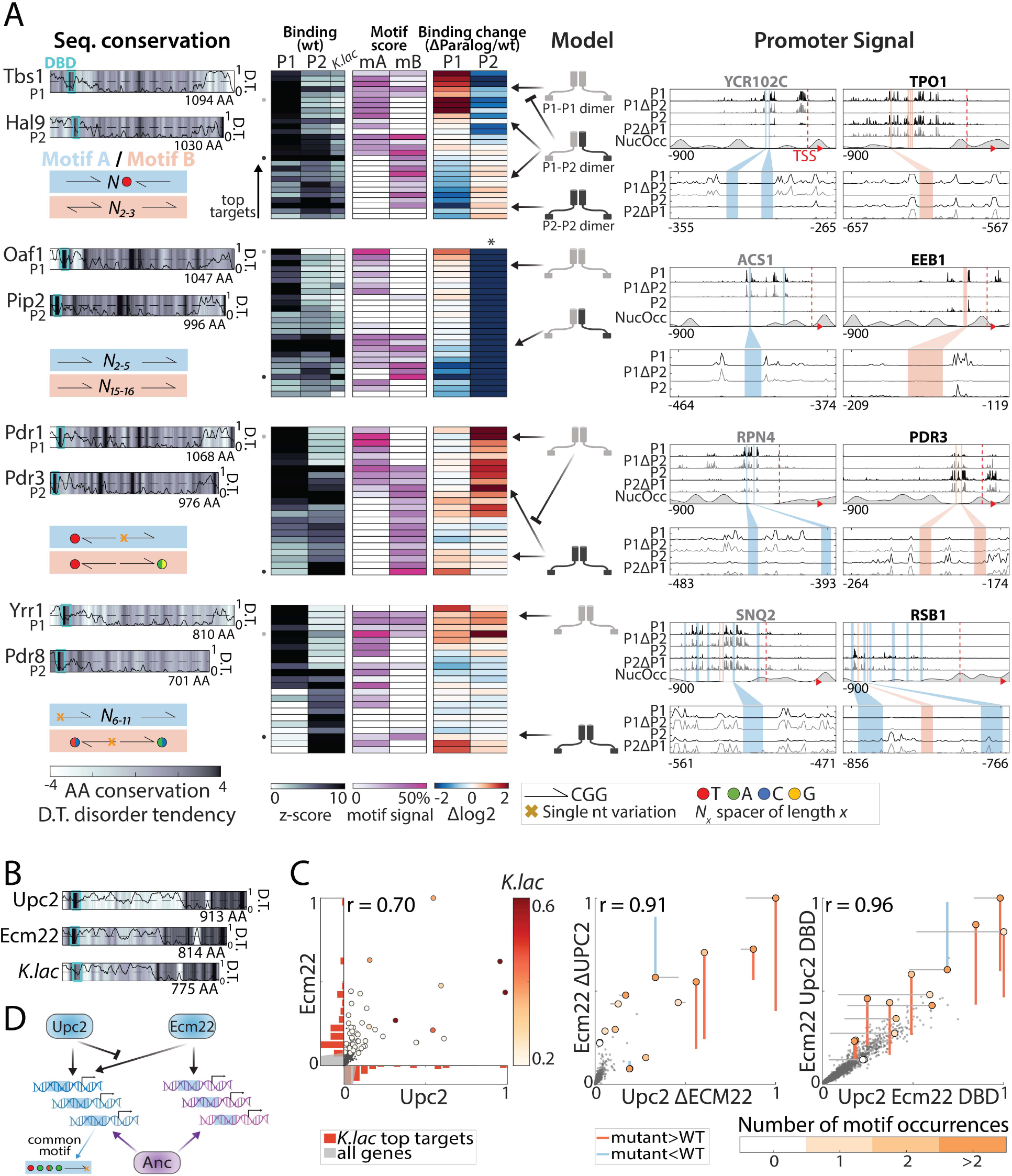
Divergence of zinc-cluster TF paralogs correlates with changes in motif preferences. **(A)** Dimerization and changes in motif-preferences may explain divergence of zinc-cluster paralogs: Zinc-cluster paralogs vary in sequence and localized at different variants of their characteristic motif, shown on the left (AA sequence conservation in color-code, DBD indicated as cyan box, and disorder tendency (*76*) shown as black line; motif symbols indicated on the bottom, see fig. S5 for motif sequences). For each pair, top-bound promoters were selected, and peak-proximal motifs defined. Shown, as indicated, are promoter binding (z-score), percentage of total promoter signal 50 bases around the indicated motifs, and binding change upon paralog deletion (log2). Suggested models explaining divergence and signal on exemplary promoters (indicated by small dots next to the binding panel) are also shown. **(B-D)** Upc2/Ecm22 diverge through DNA-binding competition: shown in (B) is the disorder tendency and pair-wise sequence conservation of Upc2-Ecm22 and the *K. lactis* ortholog with Upc2. Promoter binding preferences in the indicated backgrounds are shown in (C). Large dots indicate top 50 *K. lactis* targets, color-coded by binding strength. Distribution of these targets across the Upc2/Ecm22 binding preference are shown as histograms (red, grey: all genes). Note that Upc2 and Ecm22 bind comparably to strong *K. lactis* targets, while Ecm22 dominates on the low-intermediate targets. Colors in mid/right panels indicate the number of occurrences of the known *in-vitro* motif (TA(T/A)ACGA) and lines show change in binding relative to the wild type. A suggested model summarizing these results is depicted in (D).

We searched for CGG-containing motifs in regions bound by the zinc-cluster TFs in our dataset. In three of the five diverging pairs, differences in binding preferences correlated with differential motif preferences (Fig. 5A: differential spacing/orientation - Hal9/Tbs1 and Oaf1/Pip2 or variation within the CGG - Pdr1/Pdr3). All three pairs interacted: Hal9/Tbs1 recruited each other to their preferred sites, likely through heterodimerization, and Pip2 localized exclusively to a subset of Oaf1-preferred targets in an Oaf1-dependent manner, consistent with obligatory heterodimerization (*70*). Notably, binding preferences of both heterodimers correlated well with that of the *K. lactis* ortholog. In the case of Pdr1/3, interaction took the form of competition, with Pdr1 outcompeting Pdr3 from accessing its preferred motif.

In these three cases described above, binding preferences evolved through a combination of DNA-motif preferences and protein-protein interactions. Contrasting these, CGG-containing motifs at Yrr1/Pdr8- or Upc2/Ecm22-bound sites did not distinguish between paralogs (Fig. 5A,C and fig. S5). Rather, paralogs localized to the same motifs, but within different promoters. This divergence of promoter preferences was largely DBD-independent in the case of Yrr1/Pdr8, but greatly benefitted from DBD-dependent competition in the case of Upc2/Ecm22 (Fig. 5C). In fact, upon UPC2 deletion or DBD swapping, Ecm22 gained access to strong Upc2 sites (Fig. 5C; note correspondence to *K. lactis* sites). Of note, while DBD-swapping retained binding at the TF-specific sites, it also allowed increased access to non-specific sites, suggesting co-evolution of the DBD and the linker domain, both of which varied substantially between the paralogs. We conclude that zinc-cluster paralogs evolved largely, but not exclusively, through changes in motif preferences or affinity, resulting from combined effects of variations within, and outside, their DBDs.

### Resolution of paralog interference through competitive binding

Our analysis above described plausible evolutionary scenarios governing zinc-cluster paralog divergence, where motif preferences appear to play an important role. In most other paralogs, changes in motif preference, if exist, appear secondary to the major effective variations located outside the DBD. Still, even in such cases, DBD variations may play a role in resolving paralog interference (*45, 71*). Thus, since the two paralogs are initially redundant, divergence of binding preferences requires not only the acquisition of differential preferences, but also limiting residual, and possibly interfering paralog’s cross binding. In the case of the non-WGD pair Mcm1/Arg80, for example, such interference was resolved by weakening direct Arg80-DNA binding (*45, 71*). We asked whether similar effects can be observed for WGD paralogs.

DBD variations contributed little to the divergence of Rph1/Gis1, the most diverged C2H2 zinc-finger paralogs. We noted, however, that Rph1 gained residual access to Gis1-binding sites upon GIS1 deletion or DBD swapping (Fig. 6A,B), and Rph1-exclusive targets contained a specific variant of the common Gis1/Rph1 motif, lacking from Gis1 binding sites (Fig. 6C,D). Phylogenetic analysis revealed two variations between the paralogs: first, Gis1 has lost a conserved demethylase activity (*72*), an event that occurred quite early following WGD (Fig. 6E and fig. S6). Second, Rph1’s DBD experienced a more recent (minor) variation altering a conserved residue. This suggests the following evolutionary scenario: first, a loss of demethylase activity allowed Gis1 to specialize towards a subset of (weaker) ancestral targets and acquire new sites, through a primarily DBD-independent evolution. Second, mutations within Rph1’s DBD prevented its binding to Gis1-specialized sites, thereby reducing paralog interference (Fig. 6E).

**Fig. 6.**
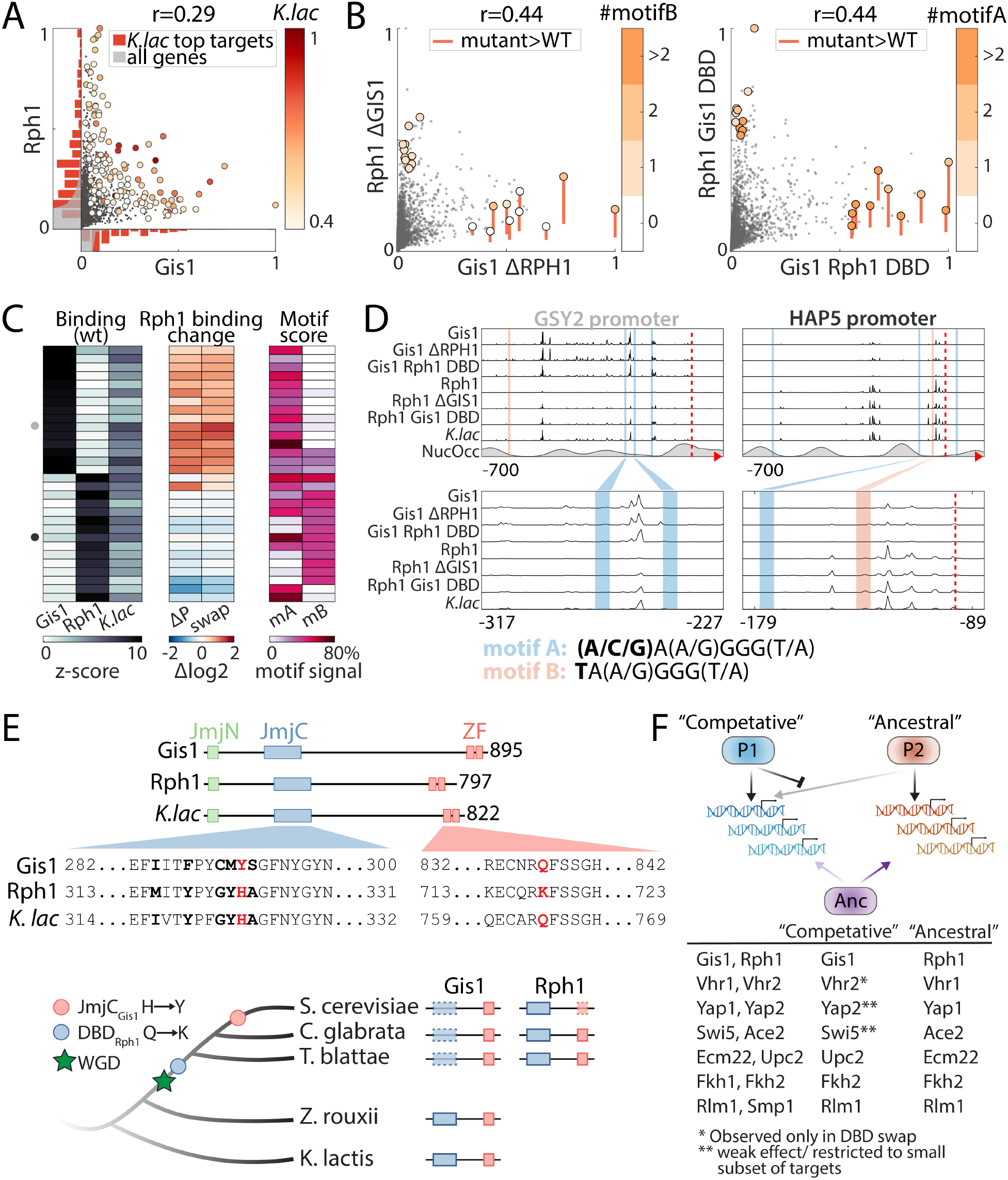
Resolution of paralog interference through competitive binding. **(A-D)** Gis1 limits Rph1 binding through DBD-dependent binding competition: shown are promoter binding preferences of Gis1 and Rph1 in wild type backgrounds (A, as in Fig. 5C) and following mutual paralog-deletion and DBD swapping (B, colored by the number of occurrences of the two known motif variants specified in D). The analysis of all top-bound promoters is summarized in (C) and binding signals on exemplary promoters are shown (D, as described in Fig. 5A). Note the increased binding of Rph1 to Gis1 target promoters upon GIS1 deletion or DBD swapping (e.g. GSY2), and reduced Gis1 binding to its target promoter after DBD swapping (e.g. HAP5). **(E)** Gis1’s loss of demethylase activity preceded variation in Rph1’s DBD: The conserved JmjC domain providing Rph1 a histone demethylase activity is mutated in Gis1 orthologs of all post-WGD species. The respective DBDs differ in only four positions, one of which replaced a conserved glutamine by lysine specifically in Rph1 and its closest orthologs (fig. S6). This suggests that the divergence was triggered by the loss of demethylase function and DBD-independent acquisition of new targets by Gis1, and finally mutation in Rph1-DBD to reduce residual Rph1 binding interference at the newly acquired Gis1 sites. **(F)** Resolution of paralog interference among diverging TF paralogs: A model for the resolution of paralog interference through competitive binding. The TF inhibiting its paralog’s binding is denoted as ‘competitive’, while the TF whose binding preference better resembles that of the *K. lactis* ortholog is denoted as ‘ancestral’. In addition to Gis1/Rph1, other diverging paralogs whose K. *lactis* orthologs were profiled appear to conform to this general model (fig. S6). Note that in most cases (indicated), divergence in promoter binding is driven by variations outside the DBD, with competition only refining, but not driving this divergence of target preferences.

Finally, extending the analysis to other diverging paralogs suggested additional cases conforming to this same design showing limited binding competition (Fig. 6F and fig. S6). Together, these results suggest a common path, whereby DBD-independent divergence is complemented by asymmetric competition, limiting residual paralog interference.

## Discussion

The binding of TFs at individual regulatory regions can vary through mutations that alter the DNA sequence or mutations that change TF binding preferences. While promoter mutations are gene specific, changes in TF binding preferences will affect multiple genes, and are therefore less likely to occur (*15, 66, 67*). Evolution of binding preferences, however, is possible through TF duplication, which generates two redundant copies and relaxes selection. The evolutionary paths through which TF paralogs diverge their binding preferences may therefore hold the key to understanding the principles through which TFs select their binding locations across the genomes.

Studying a comprehensive set of WGD-retained TF paralogs, we found that the majority of pairs still share a large fraction of overlapping targets. In fact, even diverged paralogs still localized to common targets, although at different relative strengths. This gradual divergence was not explained by variations in the DBDs. In fact, we presented evidence that, with the exception of the zinc-cluster family, variations within the DBD contributed little to binding divergence. DBD preferences may play a minor role in setting promoter binding preferences but are primarily important for stabilizing binding within selected promoters. Further, cooperation and competition act to adjust binding profiles and limit paralog interference but, with few exceptions, are not the major factors guiding divergence. In this context, gradual, and promoter-specific divergence is harder to explain within prevailing models for TF specificity, but may be more consistent with the paradigm of distributed specificity we recently proposed (*1, 73*).

At the more global level, since duplication is the major means through which new TFs emerge, the evolutionary trajectories of retained paralogs shape the transcriptional network’s design. Duplicates that diverge through sub-functionalization, for example, will confer a hierarchical design, where regulatory modules are gradually refined as the network expands. By contrast, neofunctionalization may support a distributed design, where new regulatory modules can emerge largely independent of previous connectivity. Our results reveal that neo-functionalization is quite common, although it is often combined with a biased sub-functionalization, namely uneven splitting of ancestral targets.

Finally, we focused our analysis on diverging paralogs, but it is notable that a significant fraction of paralog pairs (~40%) still binds at practically identical sites. Retention of these paralogs is therefore due to other properties. Duplicates’ tendency to cross-bind their own promoters may suggest that interactions between duplicates have evolved to confer beneficial dynamic properties not achieved by a single TF (*15, 74, 75*). Further studies are required to examine the potential advantages provided by such circuit-forming duplicates.

## Acknowledgments

We like to thank Sagie Brodsky, Tamar Jana and Offir Lupo for their strains and all Barkai lab members for fruitful scientific discussions and careful reading of the manuscript.

## Author contributions

Conceptualization: NB, TG; Methodology: TG, FJ, NB; Investigation: TG, RM; Visualization: TG, FJ; Formal analysis: TG, FJ; Data curation and Software: TG, FJ; Funding acquisition: NB; Supervision: NB; Writing – original draft: NB, TG, FJ.

## Data and materials availability

All yeast strains are available upon request, raw and processed data is available at GEO (GSE179430). MATLAB scripts for analysis and visualization are available upon request.

## Supplementary Materials

Materials and Methods

Figs. S1 to S6

Tables S1 to S4

References (*77–100*)

