## Supplementary Materials for "Evolution of binding preferences among whole-genome duplicated transcription factors"

**This PDF file includes:**

Materials and Methods

Figs. S1 to S6

Captions for Tables S1 to S4

Supplementary references

**Other Supplementary Materials for this manuscript include the following:**

Table S1 to S4 (in excel format)

Table S1. WGD-generated TF paralog pair selection

Table S2. Top targets for each TF paralog pair (compare Fig. 1F)

Table S3. Yeast strains used in this study

Table S4. Primers used to prepare DBD-swap and *K. lactis* strains

### Materials and Methods

#### Strains and Constructs

Plasmids:

All CRISPR transformations were performed using the bRA89 backbone plasmid (*1*), encoding Cas9, the target-specific guide-RNA and Hygromycin resistance. The target-specific spacer RNA template was designed using CHOPCHOP (*2*) and ligated into the pre-cut bRA89 vector as previously described (*1*) and transformed into *E .coli* for propagation. Plasmids were verified with PCR and purified with MiniPrep Kit (Real Genomics).

Yeast:

All genetic manipulations were performed in the *S. cerevisiae* BY4741 background (*3*), with the MATa his3Δ1 leu2Δ0 met15Δ0 ura3Δ0 genotype. Transformations were performed using the LiAc/SS DNA/PEG method (*4*). Following validation, the bRA89 plasmid from positive colonies was lost by growth in YPD (yeast extract peptone dextrose) and selection for colonies without bRA89-encoded Hygromycin resistance. Specific genotypes of all strains used in this study are listed in table S3.

Wild type TFs tagged with micrococcal nuclease (MNase):

TF open reading frames (ORFs) were either C- or N-terminally tagged with MNase using the C-/N-terminal SWAp-Tag (C-SWAT, N-SWAT) libraries (*5, 6*) as parental strains. The SWAT acceptor module was replaced with MNase using CRISPR. Yeast cells were transformed with a repair template (PCR amplified MNase CDS from the pGZ108 plasmid (*7*) and bRA89 plasmid with guide-RNA targeting the SWAT acceptor module. Colonies were confirmed using PCR and gel electrophoresis followed by DNA sequencing. Few strains were provided by other members of the lab (*8, 9*).

DBD Swapping strains:

Swapping strains were generated from the wild type, MNase-tagged TF background strains, using CRISPR. The cells were transformed with a genomic PCR amplification product of the corresponding paralog's DBD sequence as repair template and a locus-specific bRA89 plasmid. Colonies were confirmed using PCR and gel electrophoresis followed by DNA sequencing. Used DBD annotations are shown in fig. S2 and primers used to prepare strains are listed in table S4.

Paralog deletion strains:

Deletion strains were generated from the wild type MNase-tagged TF background strains using homologous recombination of a PCR-amplified Kanamycin or Nourseothricin resistance cassette from the pBS7 (*10*) or pFA6natNT2 (*11*) plasmid, respectively. Colonies were confirmed using PCR and gel electrophoresis.

*K. lactis* ortholog gene replacement:

*K. lactis* ortholog replacement strains were generated from the deletion strains, using CRISPR. The cells were transformed with a *K. lactis* genomic PCR amplification product of the corresponding ortholog gene sequence as a repair template and locus-specific bRA89 plasmid. The *K. lactis* gene was inserted to replace the MNase-tagged TF ORF, keeping the endogenous promoter and the MNase tag. Colonies were confirmed using PCR and gel electrophoresis followed by DNA sequencing. Primers used to prepare strains are listed in table S4.

ChEC-seq experiments:

The experiments were performed as described previously (*7*), with some modifications. Yeast strains were freshly thawed before experiments from a frozen stock, plated on YPD plates, and grown. Single colonies were picked and grown overnight at 30°C in liquid SD (synthetic complete with dextrose) medium to stationary phase. Then, the cultures were diluted ~2x$10^3$ fold into 5mL fresh SD media and grown overnight to reach an $OD_{600}$ of 4 the following morning. Cultures were pelleted at 1,500 g for 2 minutes and resuspended in 0.5mL buffer A (15mM Tris pH 7.5, 80mM KCl, 0.1mM EGTA, 0.2mM spermine, 0.5mM spermidine, 1x cOmplete EDTA-free protease inhibitors (Roche, 1 tablet per 50mL buffer), 1mM PMSF) and then transferred to 1mL 96-well plates (Thermo Scientific). Cells were washed twice in 1mL Buffer A. Next, the cells were resuspended in 150uL Buffer A containing 0.1% digitonin, transferred to an Eppendorf 96-well plate (Eppendorf 951020401) and incubated at 30°C for 5 minutes for permeabilization. Next, we added $CaCl_2$ to a final concentration of 2mM and incubated for exactly 30 seconds to activate the MNase. The MNase treatment was stopped by adding an equal volume of stop buffer (400mM NaCl, 20mM EDTA, 4mM EGTA and 1% SDS) to the cell suspension. After this, the cells were treated with Proteinase K (20mg/ml) at 55°C for 30 minutes. An equal volume of Phenol-Chloroform pH=8 (Sigma-Aldrich) was added to extract DNA. After phenol chloroform extraction of nucleic acids, the DNA was precipitated with 2.5 volumes of cold 96% EtOH, 45mg Glycoblue and 20mM sodium acetate at -80°C for > 1 hour. DNA was centrifuged (17,000 g, 4°C for 10 min), supernatant removed and the DNA pellet washed with 70% EtOH. DNA pellets were dried and resuspended in 30uL RNase A solution (0.33mg/ml RNase A in Tris-EDTA (TE) buffer (10mM Tris and 1mM EDTA)) and treated at 37°C for 20 minutes. In order to enrich for small DNA fragments, DNA cleanup was performed using SPRI beads (Ampure XP, Beckman Coulter).

Reverse SPRI cleanup was performed by adding 0.8x (24uL) SPRI beads followed by 5 minutes incubation at RT. Supernatant was collected and the remaining small DNA fragments purified by adding additional 1x (30uL) SPRI beads and 5.4x (162uL) isopropanol, and incubating 5 minutes at RT. The beads were washed twice with 85% EtOH and small fragments were eluted in 30uL of 0.1x TE buffer.

Next Generation Sequencing (NGS) library preparation:

Library preparation was performed as described in (*12*) with slight modifications. DNA fragments after RNase treatment and reverse SPRI cleanup were used as an input to End-repair and A-tailing reaction (ERA). For each sample 20uL ERA reaction (1x T4 DNA ligase buffer (NEB), 0.5mM dNTPs, 0.25mM ATP, 2.75% PEG 4000, 6U T4 PNK (NEB), 0.5U T4 DNA Polymerase (Thermo Scientific) and 0.5U Taq DNA polymerase (Bioline)) was prepared and incubated for 20 minutes at 12°C, 15 minutes at 37°C and 45 minutes at 58°C in a thermocycler.

After ERA reaction, reverse SPRI cleanup was performed by adding 0.5x (10uL) SPRI beads (Ampure XP, Beckman Coulter). Supernatant was collected and remaining small DNA fragments purified with additional 1.3x (26uL) SPRI beads and 5.4x (108uL) isopropanol. After washing with 85% EtOH, small fragments were eluted in 17uL of 0.1x TE buffer. 16.4uL elution were taken into 40uL ligation reaction (1x Quick ligase buffer (NEB), 4000U Quick ligase (NEB) and 6.4nM Y-shaped barcode adaptors with T-overhang (*13*)) and incubated for 15 minutes at 20°C in thermocycler.

After incubation, ligation reaction was cleaned by performing a double SPRI cleanup: first, a regular 1.2x (48uL) SPRI cleanup was performed and eluted in 30uL 0.1x TE buffer. Then and instead of separating the beads, an additional SPRI cleanup was performed by adding 1.3x (39uL) HXN buffer (2.5M NaCl, 20% PEG 8000) and final elution in 24uL 0.1x TE buffer. 23uL elution were taken into 50uL enrichment PCR reaction (1x Kappa HIFI (Roche), 0.32uM barcoded Fwd primer and 0.32uM barcoded Rev primer (*13*)) and incubated for 45 seconds in 98°C, 16 cycles of 15 seconds in 98°C and 15 seconds in 60°C, and a final elongation step of 1 minute at 72°C in a thermos cycler.

The final libraries were cleaned by a regular 1.1x (55uL) SPRI cleanup and eluted in 15uL 0.1x TE buffer. Library concentration and size distribution were quantified by Qubit (Thermo Scientific) and TapeStation (Agilent), respectively. For multiplexed NGS sequencing, all barcoded libraries were pooled in equal amounts, the final pool diluted to 2 nM and sequenced on NextSeq 500 (Illumina) or NovaSeq 6000 (Illumina). Sequence parameters were, Read1: 51 nucleotides (nt), Index1: 8 nt, Index2: 8 nt, Read2: 51 nt, for NovaSeq or Read1: 38 nt, Read2: 37 nt for NextSeq.

<u>NGS data processing:</u>

Raw reads from ChEC-seq libraries were demultiplexed using bcl2fastq (Illumina), and adaptor dimers and short reads were filtered out using cutadapt (*14*) with parameters: "--O 10 –pair-filter=any –max-n 0.8 –action=mask". Filtered reads were subsequently aligned to the *S. cerevisiae* genome R64-1-1 using Bowtie 2 with the options "--end-to-end --trim-to 40 --very-sensitive". The genome coverage of fully aligned read pairs was calculated with GenomeCoverage fromBEDTools (*15*) using the parameters"-d -5 –fs 1" resulting in the position of the fragment ends, which correspond to the actual MNase cutting sites. All further processing of samples with more than 200,000 concordantly aligned reads or with >0.9 correlation among biological repeats was performed using MATLAB. First, the total coverage was normalized so that the mean coverage on the nuclear genome was one. Good repeats were selected based on internal correlation to generate the mean profile for each strain (at least 2 repeats per strain).

**Quantification and statistical analyses**

Promoter definition:

Transcription start sites (TSS) were defined by combining publicly available TSS datasets (*16-18*). Promoter region was defined from the start codon until at least 700 bp upstream of the TSS (start codon if no TSS was available), or the closest verified ORF.

Promoter binding representation:

For comparison of the normalized binding signal on specific promoter examples as shown in Fig. 1C, 3D, 5A and 6D, signals were scaled so that the upper limit represents 50%, 50%, 20% and 40% of the maximal signal height respectively, across the genome in each sample. Region of promoters shown are as following: in Fig. 1C and 3D 700 nt upstream to the start codon and 150 nt downstream into the ORF, same for Fig. 6D but with 20 nt into the ORF. In Fig. 5A, 900 nt upstream to the start codon and 100 nt downstream into the ORF. Nucleosome occupancy was taken from (*19*) and smoothened with a Gaussian filter with STD = 25 nt.

Promoter binding quantification:

Genome-wide promoter binding was calculated by summing the normalized genome coverage over the promoter region of each gene (n=5424). For comparison between different TFs, the promoter-binding signal of each promoter by a certain TF was normalized to the highest signal.

TF choice for profiling (Fig. 1, fig. S1):

After constructing MNase strains for 78 out of 82 TF paralogs, we decided to proceed only with those pairs for which: (a) both paralogs could be successfully profiled under the conditions used and (b) both paralogs mostly bind to promoters of specific target genes. We therefore excluded Rsc3/Rsc30*, Aft1/Aft2*, Haa1/Cup2*, Itc1/YPL216W* Vid22/Env11* and Nfi1/Siz1* where at least one paralog could not be profiled reliably or does not show sequence specific TF activity (indicated by an asterisk), as well as Nhp6A/Nhp6B, which displayed no clear target preference. Reb1/Nsi1 or Ixr1/Abf2 were excluded as one paralog did not localize to promoter regions but ribosomal DNA or to the mitochondria genome, respectively (see table S1).

Significant TF-promoter binding for regulatory circuit analysis (Fig. 1E):

Significant TF-promoter binding was defined by z-score threshold at the 99%-quantile but not more than 3.5.

Visualizing binding changes in scatter plots (Fig. 5C, 6B, fig. S5 and S6):

In order to define the binding changes, the promoter signals in the mutant strains (DBD swap and paralog deletion) are adjusted so that the mean signal of the Top10 wild type promoters is the same as their mean signal in the wild type TF signal.

Relative, gene-specific binding changes upon paralog deletion or DBD swapping (Fig. 3, 5, 6):

To focus on gene-specific changes in binding signal, we assumed similar binding of top targets based on the strong binding correlation between the mutants and their corresponding wild types (see fig. S3): first, a robust linear regression (MATLAB function: robustfit) between the wild type (independent variable) and the mutant promoter binding across the 100 strongest bound promoters (or more, if z-score>2) was performed. The slope of this fit was then used to adjust the mutant promoter binding (P): P_adjusted=P_mutant/slope and the adjusted value compared to the wild type binding (P_WT): log2(P_adjusted/P_WT). Significantly changing genes were defined as genes whose relative binding change exceeded the mean of the 100 strongest bound promoters by one STD.

Pip2 and Hms2 DNA binding depends on the presence of their paralogs (Fig. 3, 5, fig. S5):

DNA binding profiles of Pip2 and Hms2 in the absence of their paralogs, Oaf1 and Skn7 respectively, could not be obtained. At least 4 biological repeats of each strain showed extremely low inter-correlations of promoter binding (Spearman's r<0.18, 0.27 respectively data not shown). In addition, none of the repeats showed similarity with the wild type strain (Pearson's r<0.25, 0.42 respectively, data not shown). Pip2 acting primarily as a heterodimer with Oaf1 is supported in the literature (*20*).

"New" targets determination for neo/sub-functionalization classification of *S. cerevisiae* paralogs (Fig. 4, fig. S4):

For *S. cerevisiae* paralogs and *K. lactis* ortholog targets were defined based on a z-score threshold of 4.5 and 3.5 respectively. For each *S. cerevisiae* paralog, the fraction of targets not included in the *K. lactis* targets were defined as "new".

7-mer binding quantification:

Each genomic position was indexed according to the 7-mer sequence surrounding it (-3 to +3) with assigning the same index to forward and reverse complement sequences (8192 indexes in total). Considering the properties of ChEC-seq (MNase cutting in the vicinity of the binding site, but protection of the actual binding site by the TF), the ChEC-seq signal, representing the actual cutting sites, was processed with a filter that subtracts the 7-nt moving average from the 21-nt moving average for each position and thereby punishes cutting sites. Negative values were set to zero and the mean binding score for each 7-mer index was calculated from the processed signal in the promoter regions (excluding ORFs).

Mean *in-vivo* signal around *in-vitro*-motif occurrences:

Position weight matrixes (PWMs) of *in-vitro* motifs (obtained by Protein Binding Microarrays (PBM)) were downloaded from CISBP (*21*). In the case of more than one available PWM, all *in-vitro* PWMs of this paralog pair were compared using correlation distance. The PWM couple with the highest correlation was chosen as the PWMs for this paralog pair. For better comparability between the TFs, only the most probable bases at the five most informative positions were used (in-between bases were replaced by N) to find occurrences in promoters. The mean signal for the 300-nt window centered on these occurrences was calculated. These simplified motifs were also used to select the *in-vitro*-motif-containing *in-vivo* 7-mers (Fig. 2).

DBDs comparison between paralogs:

For each paralog pair, DBD sequences based on Pfam DBD positions (*22*), determined using hmm-scan, were extracted and aligned using hmm-align (*23*). When the domain was part of the similarity regression (SR) analysis (*24*), the conserved amino acid residues were defined as every residue with >50% of the maximal conservation score. Specificity-conferring residues were defined as every residue with an SR score >150% of the mean SR score in this domain. For the remaining paralog pairs; Spt23/Mga2 (TIG), Rlm1/Smp1 (SRF) and Vhr1/Vhr2 (Vhr1), conservation score was obtained from Pfam domain HMMs using Skylign (*25*) and used to determine conserved amino acids residues like above. Specificity-defining residues were not determined for these paralog pairs (see Fig. 2, fig. S2).

Full length protein sequence conservation and phylogeny analysis of non-DBD sequences:

For each paralog pair, all orthologs sequences were downloaded from YGOB (*26*) and their Pfam-based DBD positions determined using hmm-scan (*23*). Full protein sequences and non-DBD sequences (after removing the DBDs) were aligned using m-coffee (*27*) with the options (-method t_coffee_msa clustalo_msa mafft_msa muscle_msa kalign_msa clustalw2_msa pcma_msa). Full protein sequences of eight non-WGD species were aligned separately as well, using the same method, to estimate the ancestor consensus sequence (MATLAB function seqconsensus, Fig. 4A, E, fig. S4). Global alignments with BLOSUM62 scoring matrix were used to align and calculate amino acid conservation scores for every position of both paralogs to their *K. lactis* or *Z. rouxii* ortholog, or the *K. lactis*/ *Z. rouxii* orthologue to the non-WGD consensus (Fig. 4A,E and fig. S4).

To construct the maximum likelihood tree from a constrained input tree, the non-DBD sequence alignments were then used as an input for IQTree (*28*) with ultra-fast bootstrapping (*29*), options: "-m JTT+I+G4+F -bb 1000 -g inputtree -wsr -asr -redo". The constrained input tree was based upon known species phylogeny (see fig. S4) and distinguished between the *K. lactis*/*Eremothcium* clade, the *Z. rouxii*/ *T. delbrueckii* clade, the *Lachancae* clade and all post-WGD paralogs. To adjust for protein-specific differences in evolution rates, all distances on the calculated tree were normalized so that the mean distance between *T. delbruecki* and *Z. rouxii* to their last common ancestor (LCA) was 1. These normalized values are presented in Fig. 4B. For visualization, the trees were subsequently simplified by removing all leaves (and branches) belonging to species other than the *Sacchormyces strictu* clade, *K. lactis, Z. rouxii* and *T. delbrueckii.*

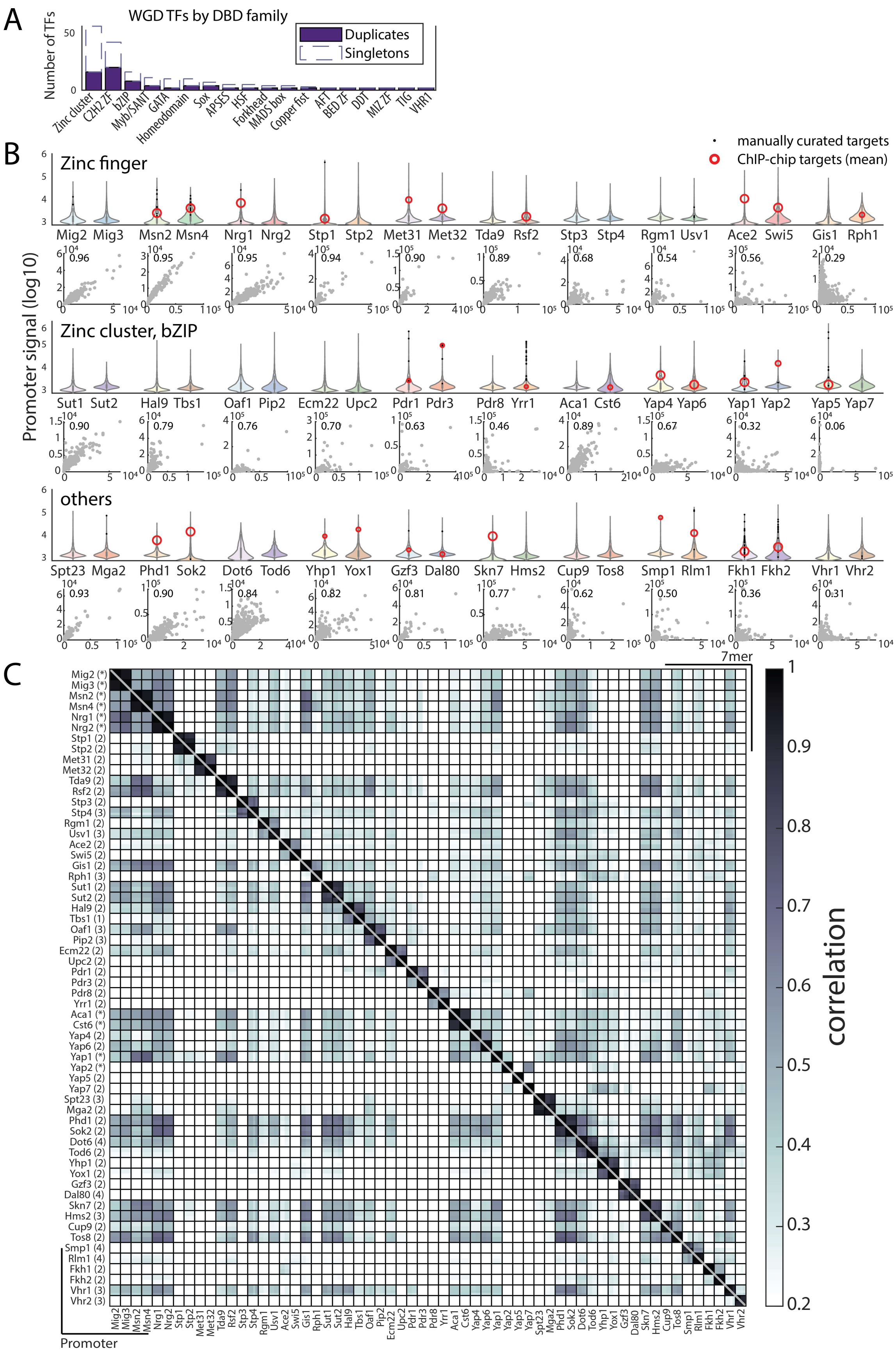

A
WGD TFs by DBD family
Number of TFs
Duplicates
Singletons
Zinc cluster
C2H2 ZF
bZIP
Myb/SANT
GATA
Homeodomain
Sox
APSES
HSF
Forkhead
MADS box
Copper fist
AFT
BED ZF
DDT
MIZ ZF
TIG
VHR1
B
manually curated targets
ChIP-chip targets (mean)
Promoter signal (log10)
Zinc finger
Mig2 Mig3 Msn2 Msn4 Nrg1 Nrg2 Stp1 Stp2 Met31 Met32 Tda9 Rsf2 Stp3 Stp4 Rgm1 Usv1 Ace2 Swi5 Gis1 Rph1
0.96 0.95 0.95 0.94 0.90 0.89 0.68 0.54 0.56 0.29
Zinc cluster, bZIP
Sut1 Sut2 Hal9 Tbs1 Oaf1 Pip2 Ecm22 Upc2 Pdr1 Pdr3 Pdr8 Yrr1 Aca1 Cst6 Yap4 Yap6 Yap1 Yap2 Yap5 Yap7
0.90 0.79 0.76 0.70 0.63 0.46 0.89 0.67 0.32 0.06
others
Spt23 Mga2 Phd1 Sok2 Dot6 Tod6 Yhp1 Yox1 Gzf3 Dal80 Skn7 Hms2 Cup9 Tos8 Smp1 Rlm1 Fkh1 Fkh2 Vhr1 Vhr2
0.93 0.90 0.84 0.82 0.81 0.77 0.62 0.50 0.36 0.31
C
7mer
Promoter
correlation
Mig2 (*)
Mig3 (*)
Msn2 (*)
Msn4 (*)
Nrg1 (*)
Nrg2 (*)
Stp1 (2)
Stp2 (2)
Met31 (2)
Met32 (2)
Tda9 (2)
Rsf2 (2)
Stp3 (2)
Stp4 (3)
Rgm1 (2)
Usv1 (3)
Ace2 (2)
Swi5 (2)
Gis1 (2)
Rph1 (3)
Sut1 (2)
Sut2 (2)
Hal9 (2)
Tbs1 (1)
Oaf1 (3)
Pip2 (3)
Ecm22 (2)
Upc2 (2)
Pdr1 (2)
Pdr3 (2)
Pdr8 (2)
Yrr1 (2)
Aca1 (*)
Cst6 (*)
Yap4 (2)
Yap6 (2)
Yap1 (*)
Yap2 (*)
Yap5 (2)
Yap7 (2)
Spt23 (3)
Mga2 (2)
Phd1 (2)
Sok2 (2)
Dot6 (4)
Tod6 (2)
Yhp1 (2)
Yox1 (2)
Gzf3 (2)
Dal80 (4)
Skn7 (2)
Hms2 (3)
Cup9 (2)
Tos8 (2)
Smp1 (4)
Rlm1 (4)
Fkh1 (2)
Fkh2 (2)
Vhr1 (3)
Vhr2 (3)
1
0.9
0.8
0.7
0.6
0.5
0.4
0.3
0.2

**Fig. S1. Sensitive, accurate and reproducible mapping of WGD TFs with ChEC-seq.**

**(A)** WGD TFs by DBD family. Shown are the number of TFs in each family divided into duplicates and singletons. **(B)** ChEC-seq identifies known and new TF target promoters. Promoter binding signal distribution for all WGD-generated TF paralogs (top Violin plot, log-scale) and direct paralog-paralog comparison (x-axis first paralog). Manually curated and Chromatin immunoprecipitation with DNA microarray (ChIP-chip) targets from SGD (*30*) are highlighted and intra-pair Pearson's correlation indicated. **(C)** Distinct and reproducible binding profiles for 60 mapped TFs. Promoter binding signal correlation (lower-left triangle) and 7-mer binding signal (upper-right triangle) correlation between all repeats for all profiled TFs (each row/column corresponds to an individual biological repeat, number of repeats indicated in parenthesis, *indicates profiles obtained from (*9*).

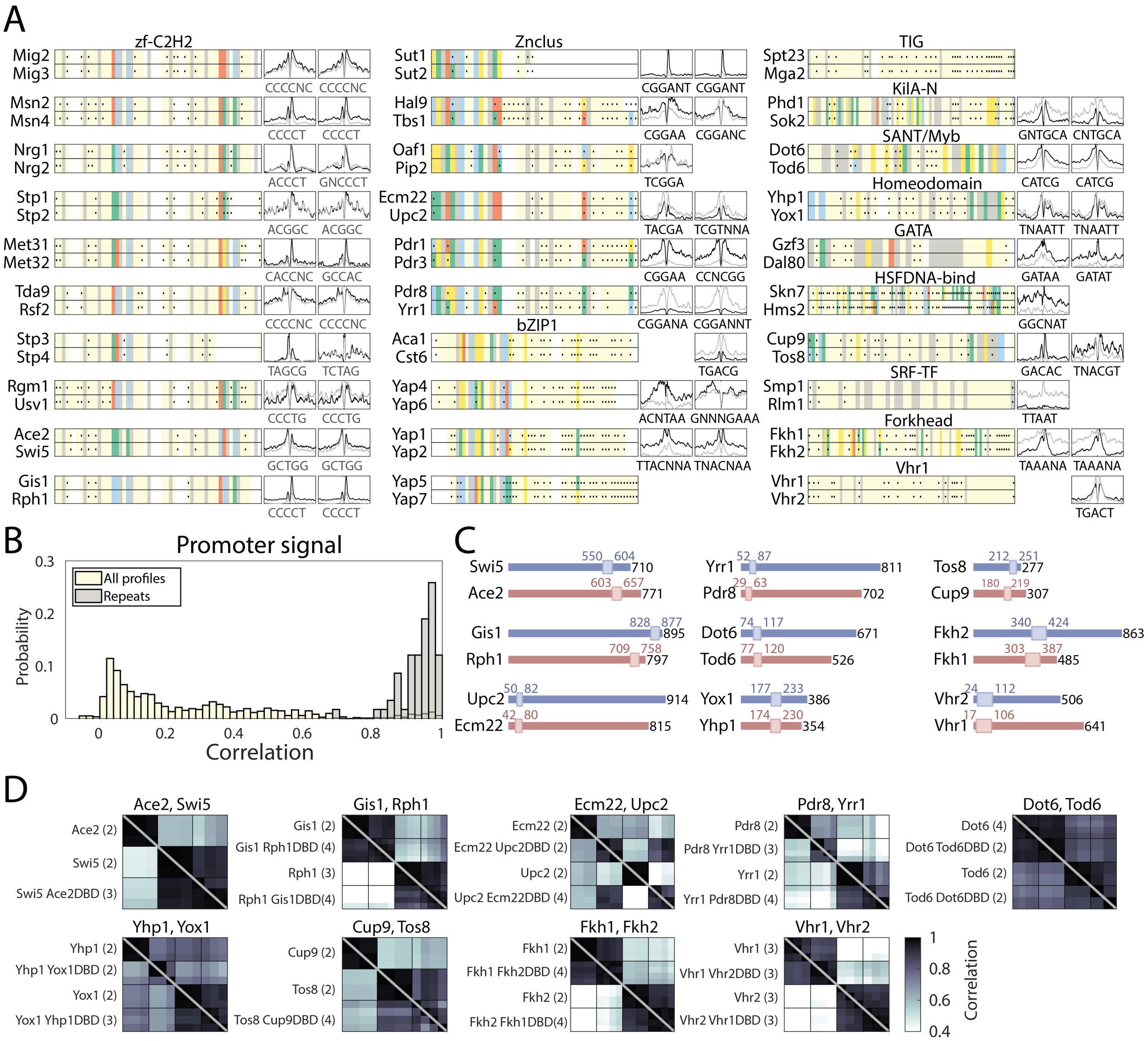

A
zf-C2H2
Mig2
Mig3
CCCCNC CCCCNC
Msn2
Msn4
CCCCT CCCCT
Nrg1
Nrg2
ACCCT GNCCCT
Stp1
Stp2
ACGGC ACGGC
Met31
Met32
CACCNC GCCAC
Tda9
Rsf2
CCCCNC CCCCNC
Stp3
Stp4
TAGCG TCTAG
Rgm1
Usv1
CCCTG CCCTG
Ace2
Swi5
GCTGG GCTGG
Gis1
Rph1
CCCCT CCCCT
Znclus
Sut1
Sut2
CGGANT CGGANT
Hal9
Tbs1
CGGAA CGGANC
Oaf1
Pip2
TCGGA
Ecm22
Upc2
TACGA TCGTNNA
Pdr1
Pdr3
CGGAA CCNCGG
Pdr8
Yrr1
CGGANA CGGANNT
bZIP1
Aca1
Cst6
TGACG
Yap4
Yap6
ACNTAA GNNNGAAA
Yap1
Yap2
TTACNNA TNACNAA
Yap5
Yap7
TIG
Spt23
Mga2
KilA-N
Phd1
Sok2
GNTGCA CNTGCA
SANT/Myb
Dot6
Tod6
CATCG CATCG
Homeodomain
Yhp1
Yox1
TNAATT TNAATT
GATA
Gzf3
Dal80
GATAA GATAT
HSFDNA-bind
Skn7
Hms2
GGCNAT
Cup9
Tos8
GACAC TNACGT
SRF-TF
Smp1
Rlm1
TTAAT
Forkhead
Fkh1
Fkh2
TAAANA TAAANA
Vhr1
Vhr1
Vhr2
TGACT
B
Promoter signal
All profiles
Repeats
Probability
0.3
0.2
0.1
0
0 0.2 0.4 0.6 0.8 1
Correlation
C
Swi5 550 604 710
Ace2 603 657 771
Gis1 828 877 895
Rph1 709 758 797
Upc2 50 82 914
Ecm22 42 80 815
Yrr1 52 87 811
Pdr8 29 63 702
Dot6 74 117 671
Tod6 77 120 526
Yox1 177 233 386
Yhp1 174 230 354
Tos8 212 251 277
Cup9 180 219 307
Fkh2 340 424 863
Fkh1 303 387 485
Vhr2 24 112 506
Vhr1 17 106 641
D
Ace2, Swi5
Ace2 (2)
Swi5 (2)
Swi5 Ace2DBD (3)
Gis1, Rph1
Gis1 (2)
Gis1 Rph1DBD (4)
Rph1 (3)
Rph1 Gis1DBD(4)
Ecm22, Upc2
Ecm22 (2)
Ecm22 Upc2DBD (2)
Upc2 (2)
Upc2 Ecm22DBD (4)
Pdr8, Yrr1
Pdr8 (2)
Pdr8 Yrr1DBD (3)
Yrr1 (2)
Yrr1 Pdr8DBD (4)
Dot6, Tod6
Dot6 (4)
Dot6 Tod6DBD (2)
Tod6 (2)
Tod6 Dot6DBD (2)
Yhp1, Yox1
Yhp1 (2)
Yhp1 Yox1DBD (2)
Yox1 (2)
Yox1 Yhp1DBD (3)
Cup9, Tos8
Cup9 (2)
Tos8 (2)
Tos8 Cup9DBD (4)
Fkh1, Fkh2
Fkh1 (2)
Fkh1 Fkh2DBD (4)
Fkh2 (2)
Fkh2 Fkh1DBD(4)
Vhr1, Vhr2
Vhr1 (3)
Vhr1 Vhr2DBD (3)
Vhr2 (3)
Vhr2 Vhr1DBD (3)
1
0.8
0.6
0.4
Correlation

**Fig. S2. Swapping experiment confirms functional conservation of DBDs between paralogs.**

**(A)** DBD sequence conservation varies between paralogs but is not associated with changes in motif selection. Shown are the aligned DBDs of both paralogs (highlighted: conserved and specificity-conferring residues) as well as their mean *in-vivo* binding signal at the known *in-vitro* motifs (compare Fig. 2A, materials and methods). **(B)** Reliable binding profiles can be obtained for DBD-swapped TFs. Distribution of promoter binding correlation between all samples (off-white) and biological repeats (grey) indicates distinct, repeatable binding profiles for DBD-swapped TFs. **(C)** DBD size and location in TFs examined in the swapping experiment. Drawn to scale are the total protein (black number indicates length in amino acids), and the position of the DBD (box and colored numbers) based on SGD Pfam annotation (red and blue correspond to the colors used in Fig. 2C,D). Shown in **(D)** are the correlations between binding preferences of the indicated TFs (individual repeats, number in parenthesis) and their swapped variants (bottom triangle promoters, top triangles 7-mers; see also Fig. 2).

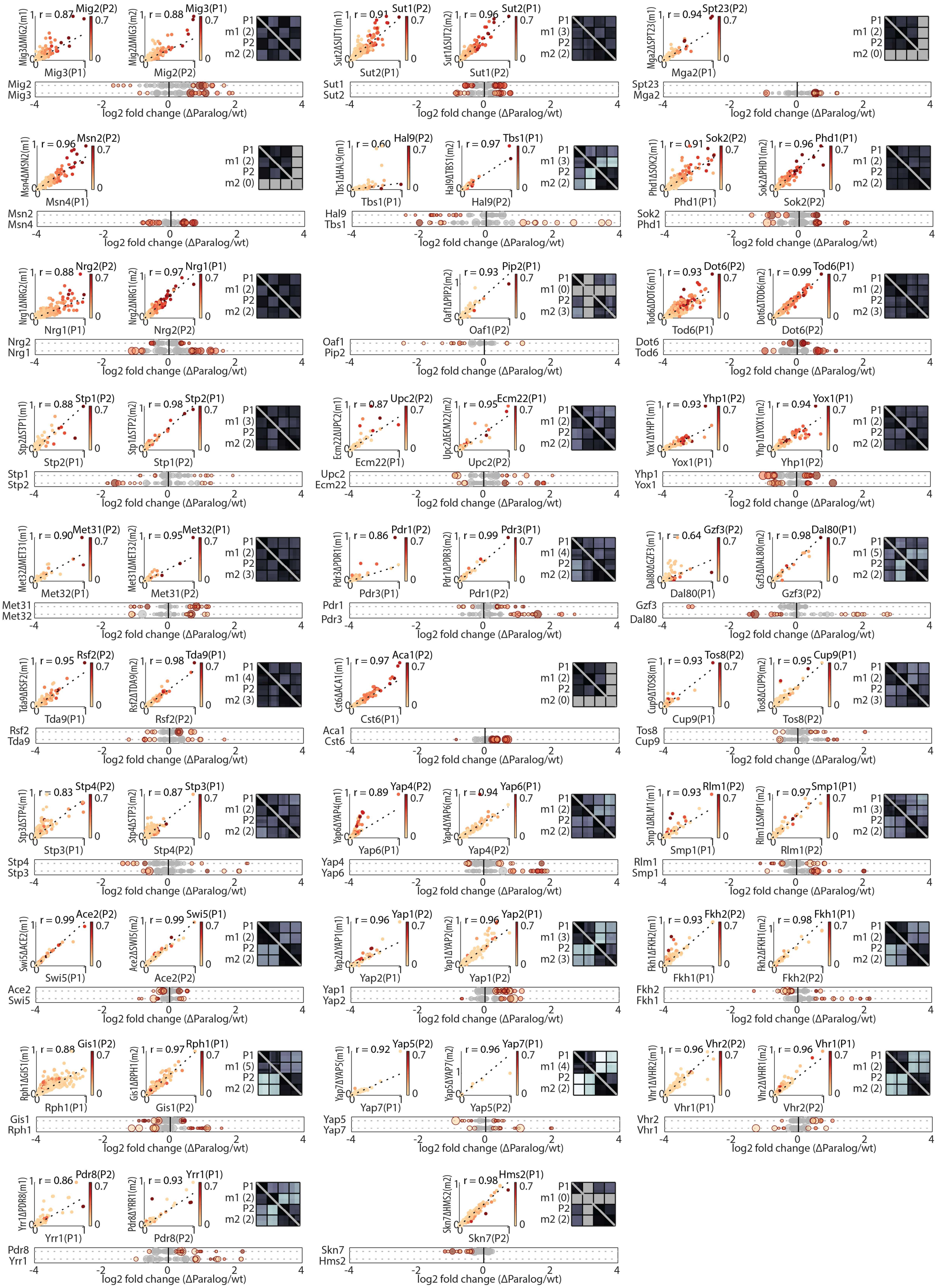

r = 0.87 Mig2(P2) Mig3ΔMIG2(m1) Mig3(P1)
r = 0.88 Mig3(P1) Mig2ΔMIG3(m2) Mig2(P2)
P1 m1 (2) P2 m2 (2)
Mig2 Mig3
r = 0.91 Sut1(P2) Sut2ΔSUT1(m1) Sut2(P1)
r = 0.96 Sut2(P1) Sut1ΔSUT2(m2) Sut1(P2)
P1 m1 (3) P2 m2 (2)
Sut1 Sut2
r = 0.94 Spt23(P2) Mga2ΔSPT23(m1) Mga2(P1)
P1 m1 (2) P2 m2 (0)
Spt23 Mga2
r = 0.96 Msn2(P2) Msn4ΔMSN2(m1) Msn4(P1)
P1 m1 (2) P2 m2 (0)
Msn2 Msn4
r = 0.60 Hal9(P2) Tbs1ΔHAL9(m1) Tbs1(P1)
r = 0.97 Tbs1(P1) Hal9ΔTBS1(m2) Hal9(P2)
P1 m1 (3) P2 m2 (2)
Hal9 Tbs1
r = 0.91 Sok2(P2) Phd1ΔSOK2(m1) Phd1(P1)
r = 0.96 Phd1(P1) Sok2ΔPHD1(m2) Sok2(P2)
P1 m1 (2) P2 m2 (2)
Sok2 Phd1
r = 0.88 Nrg2(P2) Nrg1ΔNRG2(m1) Nrg1(P1)
r = 0.97 Nrg1(P1) Nrg2ΔNRG1(m2) Nrg2(P2)
P1 m1 (2) P2 m2 (2)
Nrg2 Nrg1
r = 0.93 Pip2(P1) Oaf1ΔPIP2(m2) Oaf1(P2)
P1 m1 (0) P2 m2 (3)
Oaf1 Pip2
r = 0.93 Dot6(P2) Tod6ΔDOT6(m1) Tod6(P1)
r = 0.99 Tod6(P1) Dot6ΔTOD6(m2) Dot6(P2)
P1 m1 (2) P2 m2 (3)
Dot6 Tod6
r = 0.88 Stp1(P2) Stp2ΔSTP1(m1) Stp2(P1)
r = 0.98 Stp2(P1) Stp1ΔSTP2(m2) Stp1(P2)
P1 m1 (3) P2 m2 (2)
Stp1 Stp2
r = 0.87 Upc2(P2) Ecm22ΔUPC2(m1) Ecm22(P1)
r = 0.95 Ecm22(P1) Upc2ΔECM22(m2) Upc2(P2)
P1 m1 (2) P2 m2 (2)
Upc2 Ecm22
r = 0.93 Yhp1(P2) Yox1ΔYHP1(m1) Yox1(P1)
r = 0.94 Yox1(P1) Yhp1ΔYOX1(m2) Yhp1(P2)
P1 m1 (2) P2 m2 (2)
Yhp1 Yox1
r = 0.90 Met31(P2) Met32ΔMET31(m1) Met32(P1)
r = 0.95 Met32(P1) Met31ΔMET32(m2) Met31(P2)
P1 m1 (2) P2 m2 (3)
Met31 Met32
r = 0.86 Pdr1(P2) Pdr3ΔPDR1(m1) Pdr3(P1)
r = 0.99 Pdr3(P1) Pdr1ΔPDR3(m2) Pdr1(P2)
P1 m1 (4) P2 m2 (2)
Pdr1 Pdr3
r = 0.64 Gzf3(P2) Dal80ΔGZF3(m1) Dal80(P1)
r = 0.98 Dal80(P1) Gzf3ΔDAL80(m2) Gzf3(P2)
P1 m1 (5) P2 m2 (2)
Gzf3 Dal80
r = 0.95 Rsf2(P2) Tda9ΔRSF2(m1) Tda9(P1)
r = 0.98 Tda9(P1) Rsf2ΔTDA9(m2) Rsf2(P2)
P1 m1 (4) P2 m2 (3)
Rsf2 Tda9
r = 0.97 Aca1(P2) Cst6ΔACA1(m1) Cst6(P1)
P1 m1 (2) P2 m2 (0)
Aca1 Cst6
r = 0.93 Tos8(P2) Cup9ΔTOS8(m1) Cup9(P1)
r = 0.95 Cup9(P1) Tos8ΔCUP9(m2) Tos8(P2)
P1 m1 (2) P2 m2 (3)
Tos8 Cup9
r = 0.83 Stp4(P2) Stp3ΔSTP4(m1) Stp3(P1)
r = 0.87 Stp3(P1) Stp4ΔSTP3(m2) Stp4(P2)
P1 m1 (2) P2 m2 (2)
Stp4 Stp3
r = 0.89 Yap4(P2) Yap6ΔYAP4(m1) Yap6(P1)
r = 0.94 Yap6(P1) Yap4ΔYAP6(m2) Yap4(P2)
P1 m1 (2) P2 m2 (2)
Yap4 Yap6
r = 0.93 Rlm1(P2) Smp1ΔRLM1(m1) Smp1(P1)
r = 0.97 Smp1(P1) Rlm1ΔSMP1(m2) Rlm1(P2)
P1 m1 (3) P2 m2 (2)
Rlm1 Smp1
r = 0.99 Ace2(P2) Swi5ΔACE2(m1) Swi5(P1)
r = 0.99 Swi5(P1) Ace2ΔSWI5(m2) Ace2(P2)
P1 m1 (2) P2 m2 (2)
Ace2 Swi5
r = 0.96 Yap1(P2) Yap2ΔYAP1(m1) Yap2(P1)
r = 0.96 Yap2(P1) Yap1ΔYAP2(m2) Yap1(P2)
P1 m1 (3) P2 m2 (3)
Yap1 Yap2
r = 0.93 Fkh2(P2) Fkh1ΔFKH2(m1) Fkh1(P1)
r = 0.98 Fkh1(P1) Fkh2ΔFKH1(m2) Fkh2(P2)
P1 m1 (2) P2 m2 (2)
Fkh2 Fkh1
r = 0.88 Gis1(P2) Rph1ΔGIS1(m1) Rph1(P1)
r = 0.97 Rph1(P1) Gis1ΔRPH1(m2) Gis1(P2)
P1 m1 (5) P2 m2 (2)
Gis1 Rph1
r = 0.92 Yap5(P2) Yap7ΔYAP5(m1) Yap7(P1)
r = 0.96 Yap7(P1) Yap5ΔYAP7(m2) Yap5(P2)
P1 m1 (4) P2 m2 (2)
Yap5 Yap7
r = 0.96 Vhr2(P2) Vhr1ΔVHR2(m1) Vhr1(P1)
r = 0.96 Vhr1(P1) Vhr2ΔVHR1(m2) Vhr2(P2)
P1 m1 (2) P2 m2 (2)
Vhr2 Vhr1
r = 0.86 Pdr8(P2) Yrr1ΔPDR8(m1) Yrr1(P1)
r = 0.93 Yrr1(P1) Pdr8ΔYRR1(m2) Pdr8(P2)
P1 m1 (2) P2 m2 (2)
Pdr8 Yrr1
r = 0.98 Hms2(P1) Skn7ΔHMS2(m2) Skn7(P2)
P1 m1 (0) P2 m2 (2)
Skn7 Hms2
0 1 0.7
-4 -2 0 2 4
log2 fold change (ΔParalog/wt)

**Fig. S3. Paralog-deletion indicates gene specific paralog-paralog interactions.**

Paralog deletion only affects binding to individual genes. Shown are the direct comparison of promoter binding preferences between the wild type and mutant background, the genome wide correlation (right square) between promoter (lower-left triangle) or 7-mer binding preferences (upper-right triangle). Each row indicates a different repeat, with the total number of biological repeats indicated in parenthesis for both paralogs and their deletion mutants. Shown on the bottom is the relative binding change following paralog deletion (dot color and size indicate a TF's and its paralog's binding signal, respectively, compare Fig. 3).

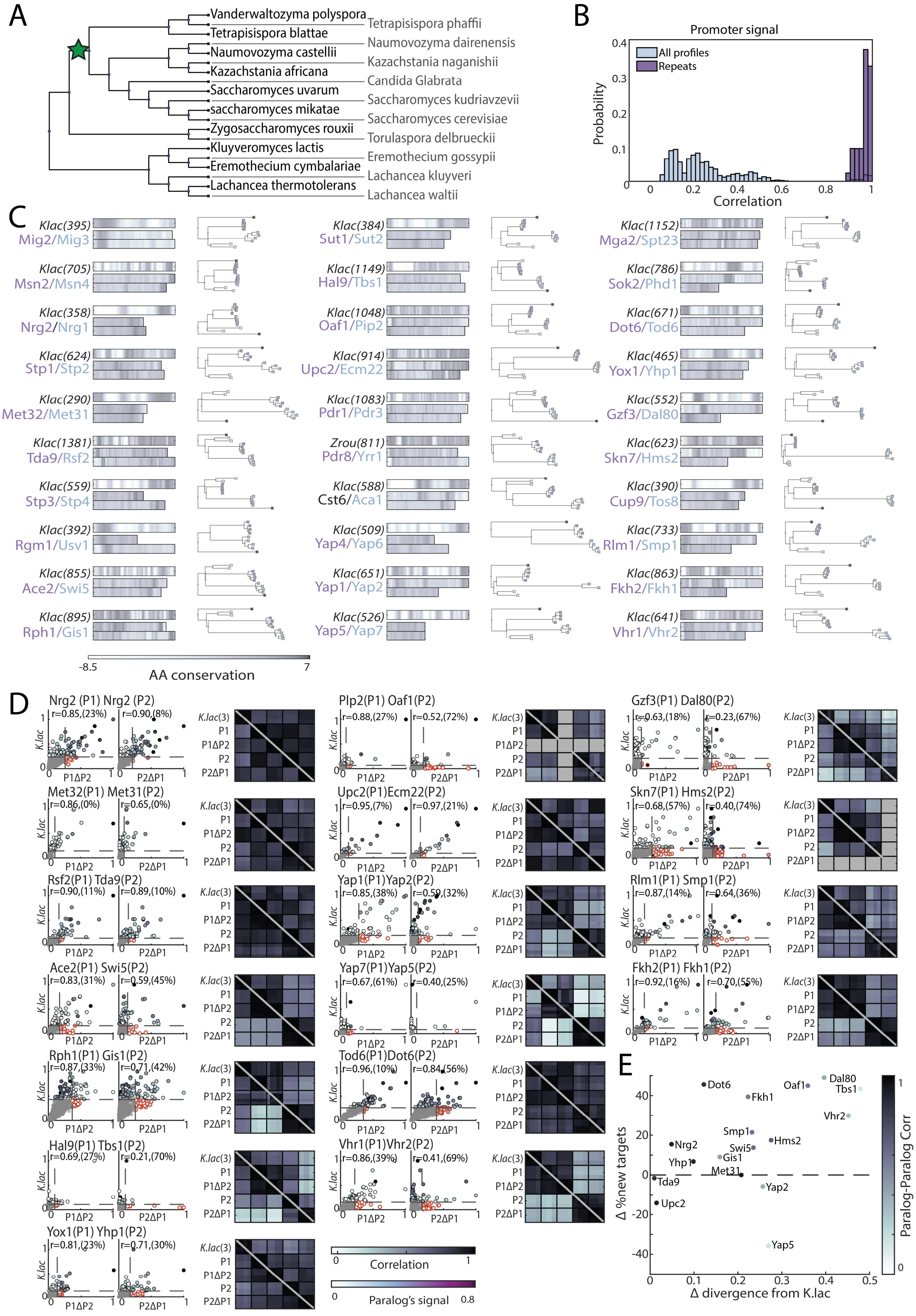
A
Vanderwaltozyma polyspora
Tetrapisispora blattae
Naumovozyma castellii
Kazachstania africana
Saccharomyces uvarum
saccharomyces mikatae
Zygosaccharomyces rouxii
Kluyveromyces lactis
Eremothecium cymbalariae
Lachancea thermotolerans
Tetrapisispora phaffii
Naumovozyma dairenensis
Kazachstania naganishii
Candida Glabrata
Saccharomyces kudriavzevii
Saccharomyces cerevisiae
Torulaspora delbrueckii
Eremothecium gossypii
Lachancea kluyveri
Lachancea waltii
B
Promoter signal
All profiles
Repeats
Probability
Correlation
C
Klac(395) Mig2/Mig3
Klac(705) Msn2/Msn4
Klac(358) Nrg2/Nrg1
Klac(624) Stp1/Stp2
Klac(290) Met32/Met31
Klac(1381) Tda9/Rsf2
Klac(559) Stp3/Stp4
Klac(392) Rgm1/Usv1
Klac(855) Ace2/Swi5
Klac(895) Rph1/Gis1
Klac(384) Sut1/Sut2
Klac(1149) Hal9/Tbs1
Klac(1048) Oaf1/Pip2
Klac(914) Upc2/Ecm22
Klac(1083) Pdr1/Pdr3
Zrou(811) Pdr8/Yrr1
Klac(588) Cst6/Aca1
Klac(509) Yap4/Yap6
Klac(651) Yap1/Yap2
Klac(526) Yap5/Yap7
Klac(1152) Mga2/Spt23
Klac(786) Sok2/Phd1
Klac(671) Dot6/Tod6
Klac(465) Yox1/Yhp1
Klac(552) Gzf3/Dal80
Klac(623) Skn7/Hms2
Klac(390) Cup9/Tos8
Klac(733) Rlm1/Smp1
Klac(863) Fkh2/Fkh1
Klac(641) Vhr1/Vhr2
-8.5 AA conservation 7
D
Nrg2 (P1) Nrg2 (P2) r=0.85,(23%) r=0.90,(8%)
Met32(P1) Met31(P2) r=0.86,(0%) r=0.65,(0%)
Rsf2(P1) Tda9(P2) r=0.90,(11%) r=0.89,(10%)
Ace2(P1) Swi5(P2) r=0.83,(31%) r=0.59,(45%)
Rph1(P1) Gis1(P2) r=0.87,(33%) r=0.71,(42%)
Hal9(P1) Tbs1(P2) r=0.69,(27%) r=0.21,(70%)
Yox1(P1) Yhp1(P2) r=0.81,(23%) r=0.71,(30%)
Plp2(P1) Oaf1(P2) r=0.88,(27%) r=0.52,(72%)
Upc2(P1)Ecm22(P2) r=0.95,(7%) r=0.97,(21%)
Yap1(P1)Yap2(P2) r=0.85,(38%) r=0.59,(32%)
Yap7(P1)Yap5(P2) r=0.67,(61%) r=0.40,(25%)
Tod6(P1)Dot6(P2) r=0.96,(10%) r=0.84,(56%)
Vhr1(P1)Vhr2(P2) r=0.86,(39%) r=0.41,(69%)
Gzf3(P1) Dal80(P2) r=0.63,(18%) r=0.23,(67%)
Skn7(P1) Hms2(P2) r=0.68,(57%) r=0.40,(74%)
Rlm1(P1) Smp1(P2) r=0.87,(14%) r=0.64,(36%)
Fkh2(P1) Fkh1(P2) r=0.92,(16%) r=0.70,(55%)
K.lac
P1ΔP2
P2ΔP1
K.lac(3)
P1
P1ΔP2
P2
P2ΔP1
Correlation
Paralog's signal
E
Δ %new targets
Δ divergence from K.lac
Paralog-Paralog Corr (mutant background)
Dot6
Fkh1
Oaf1
Dal80
Tbs1
Vhr2
Smp1
Swi5
Hms2
Nrg2
Yhp1
Gis1
Met31
Tda9
Yap2
Upc2
Yap5

**Fig. S4.** ***K. lactis*** **orthologs represent possible binding preferences of the pre-duplication ancestor.**

**(A)** Schematic representation of the phylogeny of the genus *Saccharomyces*. Genomic sequences of indicated species were used in the phylogenetic analysis to assess sequence divergence rate. **(B)** Reproducible and distinct profiles of *K. lactis* orthologs profiled in *S. cerevisiae*. Distribution of promoter binding correlations between all samples (light blue) and biological repeats (purple) of the *K. lactis* orthologs. **(C)** Sequence similarity and divergence differs between paralog pairs. For each paralog pair, we compared sequences of one non-WGD ortholog (*K. lactis* or *Z. rouxii* as indicated) against the non-WGD consensus (top left, shown is the average alignment score 20 amino acids around each residue, number in parenthesis indicates protein length) and against the sequence of both *S. cerevisiae* paralogs (bottom, color indicates the alignment score of the respective residue). Rate of non-DBD sequence divergence was derived from phylogenetic analysis of non- and post-WGD orthologs (right, reduced phylogeny is shown with dots indicating different orthologs; black: *K. lactis*, white: *Z. rouxii*, colored: paralogs of the *Saccharomyces strictu* lineage corresponding to similar colored *S. cerevisiae* proteins). **(D)** *K. lactis* orthologs reflect paralogs' binding preferences and suggest neo-functionalization as the dominant divergence principle. For each paralog pair, comparing *K. lactis* ortholog to each TF in the deletion background revealed the binding preference conservation (left, r indicates Pearson's correlation). These correlations can be modified by paralog interactions present in the wild type strains (right, each row indicates a different repeat, with the total number of repeats for *K. lactis* samples indicated in parenthesis). The fraction of "new" targets acquired by *S. cerevisiae* paralogs is indicated (in parenthesis) and corresponds to the red-outlined dots (see materials and methods). **(E)** Divergence is associated with new target acquisition in most paralog pairs. Shown is the divergence difference between both paralogs from the *K. lactis* ortholog and the difference between the size of the fraction of newly acquired targets. Note that the more diverged paralog (indicated) also acquired more new targets.

A

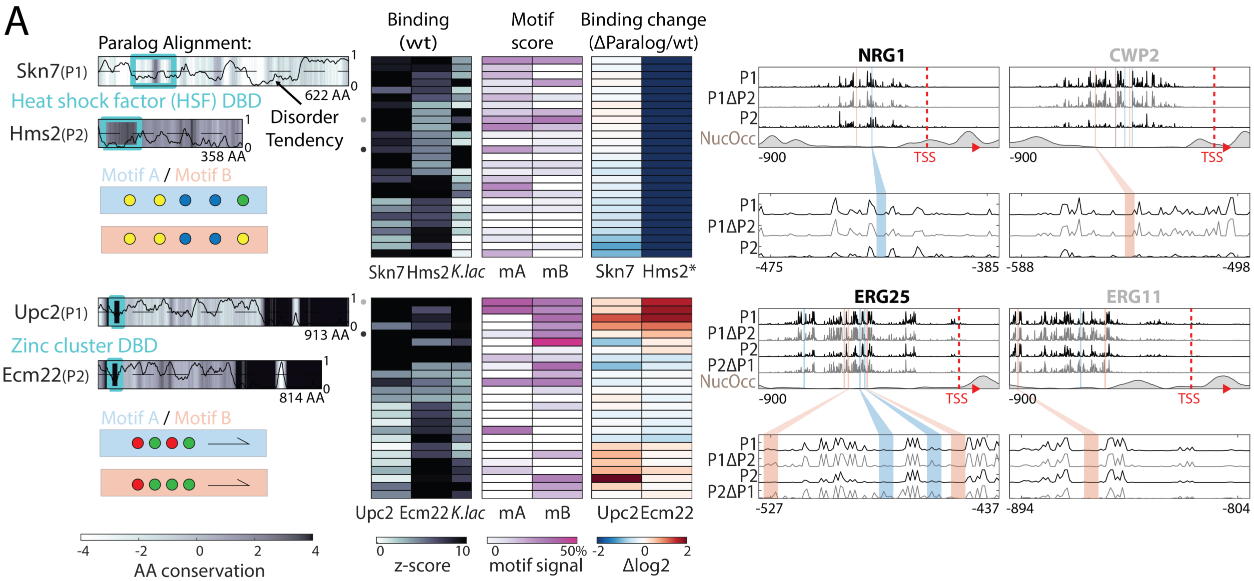


B

| *Paralog Pair* | *Motif A* | | *Motif B* | |
|---|---|---|---|---|
| **Tbs1, Hal9** | N | CGGNTCCG | $N_{2-3}$ | C(G/C)G$N_{2-3}$CGG |
| **Oaf1, Pip2** | $N_{2-5}$ | CGG$N_{2-5}$CGG | $N_{15-16}$ | CGGN CCG |
| **Pdr1, Pdr3** | | TCCGTGG | | TCCGCGG(A/G) |
| **Yrr1, Pdr8** | $N_{6-11}$ | (T/C)GG$N_{6-11}$CGG | | (T/C)CC(G/T)CGG(A/C) |
| **Upc2, Ecm22** | | TATACGA | | TAAACGA |
| **Skn7, Hms2** | | GGCCA | | GGCCG |

**Fig. S5. Homo- and heterodimerization's role in the binding preference and divergence of dimer-forming paralog pairs.**

**(A)** Heterodimer formation explains asymmetric dependency in the Skn7/Hms2 paralog pair of the HSF DBD family. Shown are the AA sequence conservation in color-code, DBD indicated as cyan box, and disorder tendency shown as black line. Motif symbols indicated on the bottom. Also shown is the binding behavior to the top promoter targets (binding scores of both TFs and their *K. lactis* ortholog, log2 change after paralog deletion and motif occurrences), and binding signal on representative promoters (compare Fig. 5A). **(B)** Competition shapes the binding profile of the Ecm22/Upc2 paralog pair (compare Fig. 5C). **(C)** Identified motifs in the top promoters of each paralog pair. Shown are the schematic and sequences for the proposed TF motifs found in the respective target genes.

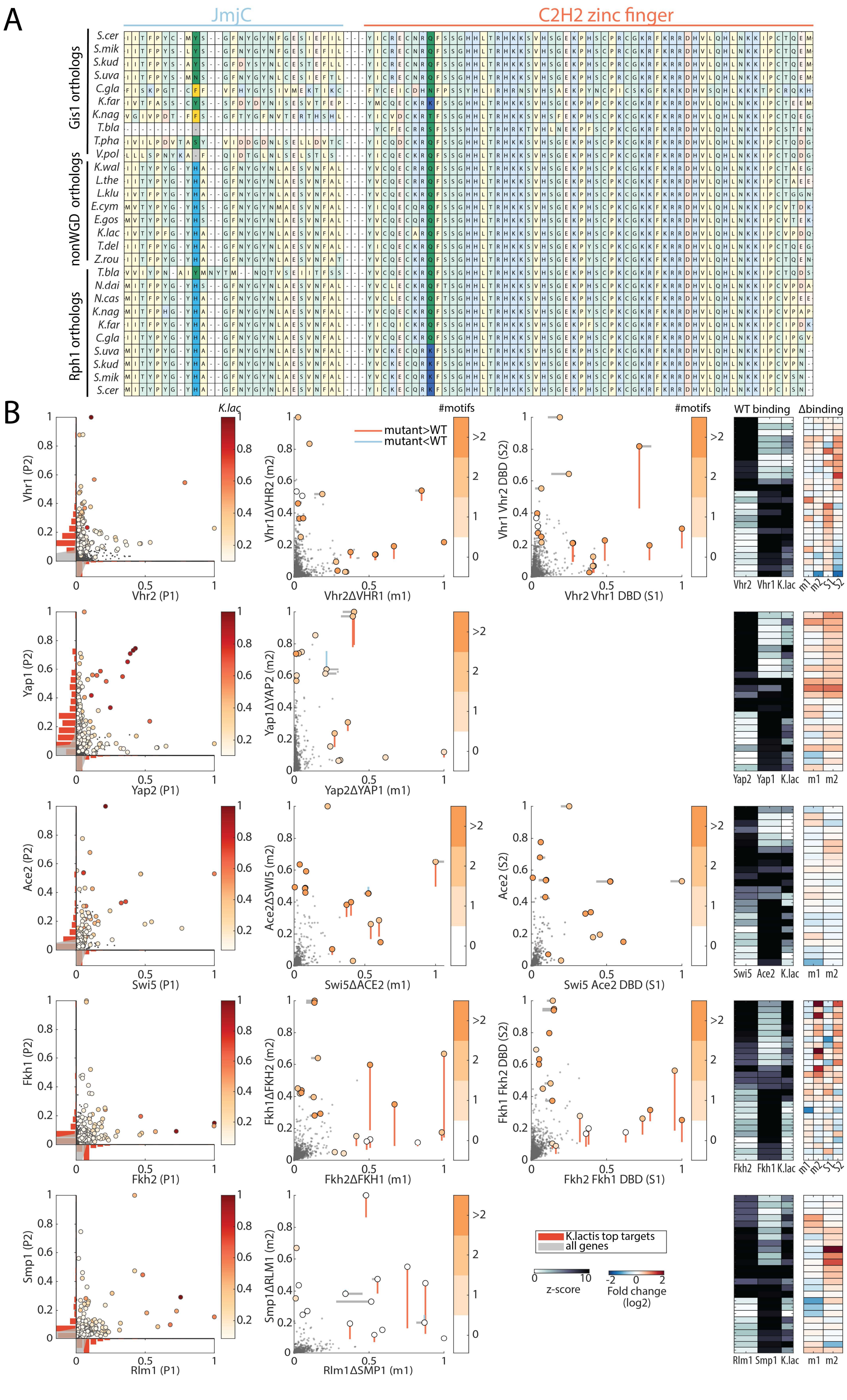
A
JmjC
C2H2 zinc finger
Gis1 orthologs
nonWGD orthologs
Rph1 orthologs
S.cer
S.mik
S.kud
S.uva
C.gla
K.far
K.nag
T.bla
T.pha
V.pol
K.wal
L.the
L.klu
E.cym
E.gos
K.lac
T.del
Z.rou
T.bla
N.dai
N.cas
K.nag
K.far
C.gla
S.uva
S.kud
S.mik
S.cer
B
K.lac
#motifs
mutant>WT
mutant<WT
#motifs
WT binding
Δbinding
Vhr1 (P2)
Vhr2 (P1)
Vhr1ΔVHR2 (m2)
Vhr2ΔVHR1 (m1)
Vhr1 Vhr2 DBD (S2)
Vhr2 Vhr1 DBD (S1)
Vhr2 Vhr1 K.lac m1 m2 S1 S2
Yap1 (P2)
Yap2 (P1)
Yap1ΔYAP2 (m2)
Yap2ΔYAP1 (m1)
Yap2 Yap1 K.lac m1 m2
Ace2 (P2)
Swi5 (P1)
Ace2ΔSWI5 (m2)
Swi5ΔACE2 (m1)
Ace2 (S2)
Swi5 Ace2 DBD (S1)
Swi5 Ace2 K.lac m1 m2
Fkh1 (P2)
Fkh2 (P1)
Fkh1ΔFKH2 (m2)
Fkh2ΔFKH1 (m1)
Fkh1 Fkh2 DBD (S2)
Fkh2 Fkh1 DBD (S1)
Fkh2 Fkh1 K.lac m1 m2 S1 S2
Smp1 (P2)
Rlm1 (P1)
Smp1ΔRLM1 (m2)
Rlm1ΔSMP1 (m1)
K.lactis top targets
all genes
z-score
Fold change (log2)
Rlm1 Smp1 K.lac m1 m2

**Fig. S6. The role of competition and paralog interference in the divergence of TF paralogs.**

**(A)** Sequence conservation of JmjC domain and DBD in the Rph1/Gis1 paralog pair. Shown is the part of the multiple sequence alignment between all Rph1/Gis1 orthologs containing the discussed changes in the JmjC domain and the first C2H2 zinc finger (highlighted in bold colors). **(B)** The impact of competition and DBD specificity on the divergence of TF paralog pairs outside the zinc cluster family. Shown are the comparisons between the promoter binding preferences of both paralogs and the *K. lactis* ortholog (left, color indicates *K. lactis* binding signal) as well as the impact of paralog-deletion (middle) and DBD swapping (right) and the quantification for the top targets, for each paralog pair discussed in Fig. 6F (m1/2: paralog deletion mutants, s1/2: DBD swapping).

**Table S1. WGD-generated TF paralog pair selection**

List of all DBD-containing WGD-generated paralogs in *S. cerevisiae* with DBD family and the performed experiments (grey font indicates filtered-out paralog pairs).

**Table S2. Top targets for each TF paralog pair (compare Fig. 1F)**

For each pair (sorted by family and inter-pair correlation) the top 40 targets based on promoter binding signal (as used in Fig. 1F) are listed with their standard name, systematic name and the promoter binding z-score in each paralog of the pair.

**Table S3. Yeast strains used in this study**

List of strains used in this study with their background genotype and source. (In the genotype-column "TF" stands for the ORF of the MNase tagged TF and "tf" for that of its paralog).

**Table S4. Primers used to prepare DBD-swap and *K. lactis* strains**

For each created DBD-swap/ortholog replacement strain the forward and reverse primers used in the genomic PCR (*S. cerevisiae* genome for DBD swaps and *K. lactis* genome for ortholog replacement, respectively) are listed.
